# Overcoming protocatechuate and catechol accumulation in muconic acid production via adaptive laboratory evolution and metabolic engineering in *Pseudomonas putida*

**DOI:** 10.64898/2026.07.14.738518

**Authors:** Alissa C. Bleem, Tracy L. Hodges, Torrey M. Lind, Eugene Kuatsjah, Yuqian Gao, Michael A. Gapuz, Zoe A. Kellermyer, Alexander F. Benson, Morgan A. Ingraham, Allison Z. Werner, Young-Mo Kim, Christopher W. Johnson, Gregg T. Beckham

**Author notes:** Molecular & Cellular Biology Graduate Program, University of Washington, Seattle, WA, USA. Institute of Sustainability for Chemicals, Energy and Environment (ISCE^2^), Agency for Science, Technology and Research (A*STAR), 138665 Singapore, Republic of Singapore. Department of Microbiology, University of Chicago, Chicago, IL, USA.

## Abstract

Muconic acid is an industrially valuable molecule that can be biologically produced from diverse biogenic and waste-derived feedstocks, including sugars and lignin- and plastic-derived aromatic compounds. However, accumulation of protocatechuate (PCA) has been observed in multiple microbes engineered for muconate production when the PCA decarboxylase, AroY, is used. This raises the question of whether PCA decarboxylation represents a rate-limiting step and how this bottleneck might be alleviated, especially given the toxicity and reactivity of PCA and catechol intermediates. To address this, we performed adaptive laboratory evolution (ALE) on a strain of *Pseudomonas putida* originally engineered for muconate production from aromatic compounds, but with *catBC* restored, to select for improved conversion of PCA and, in separate lineages, 4-hydroxybenzoate. Contrary to our expectations, the predominant beneficial mutations localized to the *catA1* cassette encoding catechol 1,2-dioxygenase, rather than *aroY* or its associated cofactor biosynthesis genes. Transcriptomic analysis revealed elevated *catA1* expression in evolved isolates from ALE, and introduction of these mutations improved productivity in strains designed for muconate production from both aromatic and sugar substrates. Quantitative proteomics and biochemical assays demonstrated that the mutations also led to increased CatA1 protein abundance and modest enhancements in catalytic efficiency, respectively, with strain phenotypes largely driven by high CatA1 levels and potentially synergistic kinetic improvements. Additional reverse-engineering studies identified variants with modest effects on muconate accumulation, including those with potential to enhance biosynthesis of the prenylated FMN cofactor of AroY. Collectively, these results indicate that catechol, not PCA, is the principal bottleneck in muconate production via the PCA decarboxylation route originally demonstrated by Draths *et al*., refining our understanding of pathway limitations and offering new strategies for improving rate, yield, and strain resilience in muconate bioproduction.

**Highlights:**

- Accumulation of metabolic intermediates was alleviated by adaptive laboratory evolution
- Sequencing, proteomics, and enzyme kinetics revealed mechanisms for adaptation
- Increased CatA1 expression reduced bottlenecks and improved muconate production

## 1. Introduction

Significant research investment has established *cis,cis-*muconic acid (hereafter, muconate) as a platform chemical that can be efficiently accessed from the catechol catabolic pathway in several microbes (Draths and Frost, 1994; Johnson et al., 2016; Wang et al., 2021; Xie et al., 2014). After separation from fermentation broth (Tönjes et al., 2023; Tönjes et al., 2024; Vardon et al., 2016), muconate can undergo catalytic conversion to direct replacement chemicals such as adipic acid, adiponitrile, and caprolactam (Beerthuis et al., 2015; Carraher et al., 2017; Draths and Frost, 1994; Huo and Shanks, 2020; Khalil et al., 2020; Müller et al., 2015; Vardon et al., 2016), or to performance-advantaged polymers (Cywar et al., 2021; Dell’Anna et al., 2021; Hu et al., 2025; Johnson et al., 2019; Luo et al., 2022; Molinari et al., 2024; Rorrer et al., 2016; Rorrer et al., 2017; Suastegui et al., 2016; Wu et al., 2025). Incumbent methods for production of these chemicals involve significant energy and greenhouse gas emissions (Nicholson et al., 2023) and bio-based approaches may offer a more sustainable approach (Corona et al., 2018; Khoo et al., 2016; Mokwatlo et al., 2024; Sudarsanam et al., 2020; van Duuren et al., 2020). Since the pioneering work of Draths and Frost for biological production of muconate from glucose in *Escherichia coli* (Draths and Frost, 1994; Frost and Draths, 1996), biological production of this compound has been demonstrated in many other bacteria and yeast, using a range of feedstocks (Almqvist et al., 2021; Cai et al., 2020; Li et al., 2024; Liu et al., 2025; Mokwatlo et al., 2024; Moon et al., 2025; Pyne et al., 2023; Vardon et al., 2015; Vilbert et al., 2024; Wang et al., 2021; Weiland et al., 2023). However, decarboxylation of protocatechuate (PCA) to catechol and ring-opening of catechol to muconate—the penultimate and terminal enzymatic steps for muconate production, respectively (**Fig. 1A**)—represent metabolic bottlenecks that can limit yield. Indeed, accumulation of PCA and/or catechol has been observed in *E. coli* (Choi et al., 2019; Molinari et al., 2024), *Pseudomonas putida* (Kuatsjah et al., 2022; Ling et al., 2022; Mokwatlo et al., 2024), *Saccharomyces cerevisiae* (Tönjes et al., 2024; Wang et al., 2020), *Novosphingobium aromaticivorans* (Vilbert et al., 2024), *Corynebacterium glutamicum* (Li et al., 2024), *Pichia occidentalis* (Pyne et al., 2023), and *Acinetobacter baylyi* (Liu et al., 2025), underscoring the ubiquity of this bottleneck across diverse hosts and processes. A longstanding question in the field is whether PCA decarboxylation constitutes a rate-limiting step, and how this bottleneck might be alleviated given the toxicity and reactivity of PCA and catechol intermediates.

**Figure 1.**
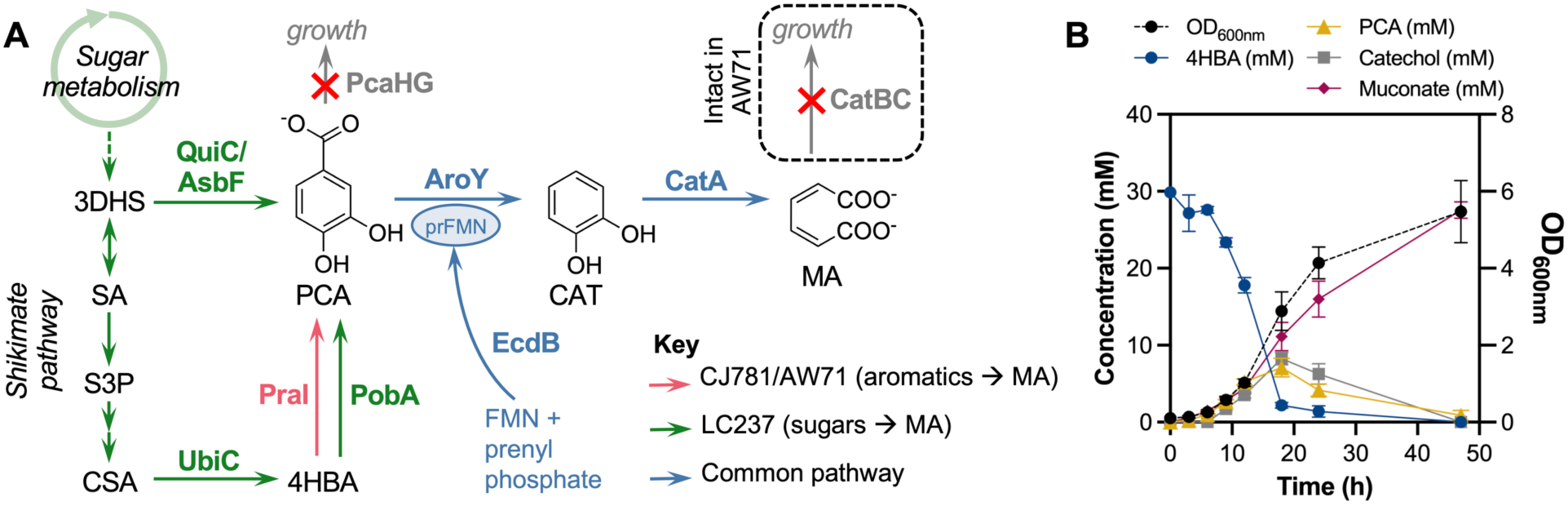
Muconate-producing strains of *P. putida* exhibit metabolic bottlenecks at PCA and catechol. **(A)** *P. putida* strains designed for muconate production from sugars convert PCA via AroY and CatA. Activity of the AroY decarboxylase requires a prenylated FMN cofactor (PrFMN), supplied by the EcdB flavin prenyltransferase. AW71, the strain used for ALE, has *catBC* intact to enable AroY-dependent growth on aromatics. **(B)** CJ781 produces PCA and catechol when cultivated in shake flasks in M9 minimal medium with 30 mM 4-hydroxybenzoate (4HBA) and 15 mM glucose. Glucose was added at 12 and 24 h to a concentration of 15 mM. Error bars indicate the standard deviation from the mean of three biological replicates. Numerical data are provided in **Supplementary File 1**. Abbreviations: 3DHS, 3-dehydroshikimate; SA, shikimate; S3P, shikimate-3-phosphate; CSA, chorismate; 4HBA, 4-hydroxybenzoate, PCA, protocatechuate; CAT, catechol; MA, muconate.

Development of an efficient biocatalyst for muconate production involves optimization of the primary production pathway as well as maintenance energy and cofactor balancing. Prior studies have demonstrated the effectiveness of *P. putida* for conversion of lignocellulosic sugars and aromatic compounds due to its rapid growth, robust metabolic tolerance, and ease of genetic manipulation (Martínez-García and de Lorenzo, 2024; Weimer et al., 2020; Wilkes et al., 2024). In prior work, we developed *P. putida* strain CJ781 (Kuatsjah et al., 2022), which uses glucose for growth while producing muconate from lignin-related aromatic compounds, and strain LC237 (Kim et al., 2026), which converts cellulosic sugars (glucose, xylose, and arabinose) to muconate in a growth-coupled manner (**Table 1**, **Fig. 1A**). Both of these strains use the AroY decarboxylase enzyme from *Enterobacter cloacae* for PCA decarboxylation to catechol, along with the EcdB flavin prenyltransferase to synthesize the prenylated flavin mononucleotide (prFMN) cofactor required for decarboxylase activity (Johnson et al., 2016; Payne et al., 2015; Sonoki et al., 2014; White et al., 2015). *P. putida* produces prFMN, but AroY apparently requires more than is produced natively (Johnson et al., 2016). The *P. putida* genome natively encodes two catechol 1,2-dioxygenases, *catA1* and *catA2*, to catalyze oxidative ring cleavage of catechol to muconate, which can be accumulated as a final product upon deletion of the downstream catabolic genes, *catBC* (**Fig. 1A**). Strong expression of *aroY*, *ecdB*, and *catA1* from the constitutive *tac* promoter was previously demonstrated to limit accumulation of metabolic intermediates (Johnson et al., 2016), but CJ781 and LC237 still accumulate significant quantities of PCA and catechol when cultivated with aromatic substrates (**Fig. 1B**, **Fig. S1)** (Kim et al., 2026). PCA and catechol are reactive molecules that exhibit toxicity to *P. putida* cells (Jiménez et al., 2014; Salvachúa et al., 2018), and even minor bottlenecks in these pathways can severely impact process economics, limiting muconate titer, rate, and overall fermentation time (Matthiesen et al., 2016; Mokwatlo et al., 2024; Werner et al., 2023).

**Table 1.**
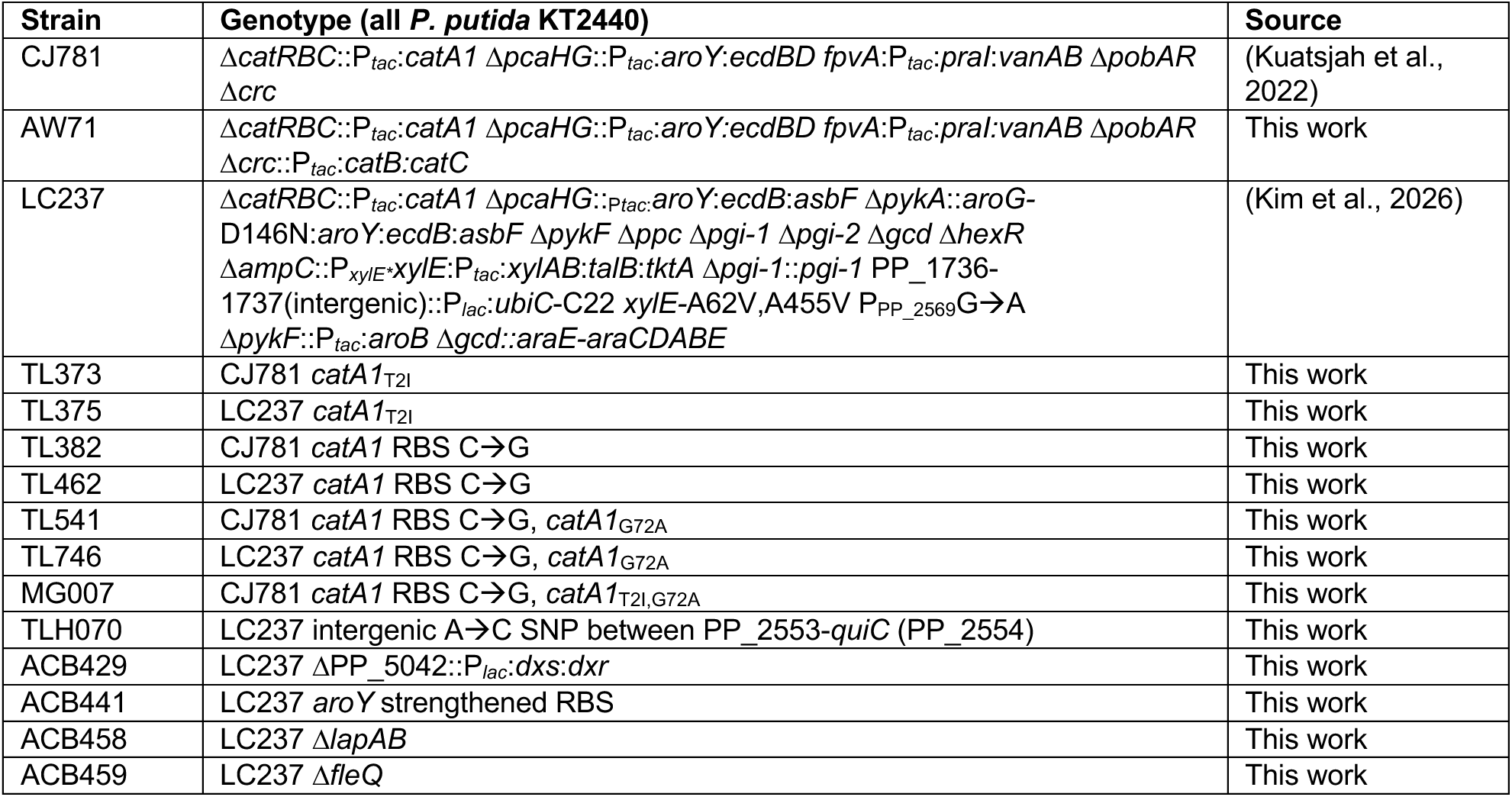
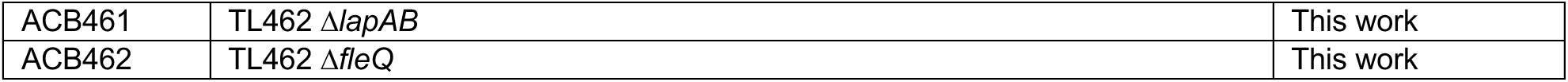
Strains used in this study.

In this work, we used adaptive laboratory evolution (ALE) with the aim to alleviate metabolic bottlenecks at PCA and catechol in strains of *P. putida* engineered for muconate production. ALE was conducted in AW71, a strain with the same genetic background as CJ781 but lacking the *catBC* gene deletions required for muconate accumulation, thereby enabling growth via PCA and catechol (**Fig. 1A**). Evolution experiments enriched for mutated strains with reduced accumulation of PCA and catechol as metabolic intermediates, and sequencing (whole-genome and transcriptomics) of the evolved mutants revealed genetic drivers of enhanced phenotypes. When muconate accumulation was restored in evolved, metabolically debottlenecked isolates, they exhibited higher muconate production rates than the unevolved parent strain from aromatic substrates. Causal mutations were also recapitulated in clean strain backgrounds for muconate production from both biomass-derived sugars and aromatics, and beneficial mutations associated with increased abundance of the CatA1 protein were shown to significantly improve the muconate production rate while eliminating accumulation of PCA and catechol as byproducts. The resulting strains represent superior biocatalysts for muconate production from sugars and aromatics.

## 2. Results

### 2.1. Evolution of an AroY-dependent strain alleviated the PCA bottleneck

To establish growth dependence on PCA catabolism during ALE, we first restored the *catBC* locus in the previously engineered CJ781 strain (Kuatsjah et al., 2022). The resulting strain, AW71 (**Fig. 1A**, **Table 1**), uses aromatic compounds as the sole source of carbon and energy while retaining all other genetic modifications as in CJ781. Like CJ781, AW71 also accumulated significant quantities of PCA and catechol during growth with 4-hydroxybenzoate (4HBA) (**Fig. 2A**), so it served as the parent strain for ALE experiments designed to overcome these bottlenecks. To interrogate PCA catabolism in AW71, two tolerance ALE (TALE) (Mohamed et al., 2020) experiments were conducted. Six independent cultures of AW71 were cultivated in minimal media containing either 15 mM 4HBA or 15 mM PCA. Each day, cells were passaged into fresh media, and the substrate was slowly increased over the course of the experiment to a final concentration of 30 mM after 44 passages (∼4,000 generations; see Methods; **Supplementary File 1**). All evolved populations exhibited improved growth on the substrate used for their selection, as evinced by decreases in growth lag times and increases in maximum specific growth rates (**Fig. 2B,C**). At the end of the experiment, evolved populations were struck out on agar plates to select a clonally unique isolate from each population, and these exhibited similarly improved phenotypes to their corresponding population samples (**Fig. S2**). Growth profiling of populations collected throughout the experiment demonstrated that the strongest improvements arose in the earlier phases of passaging, especially for lineages evolved on 4HBA, but some lineages exhibited incremental improvements throughout the experiment (**Figs. S3-S4**).

**Figure 2.**
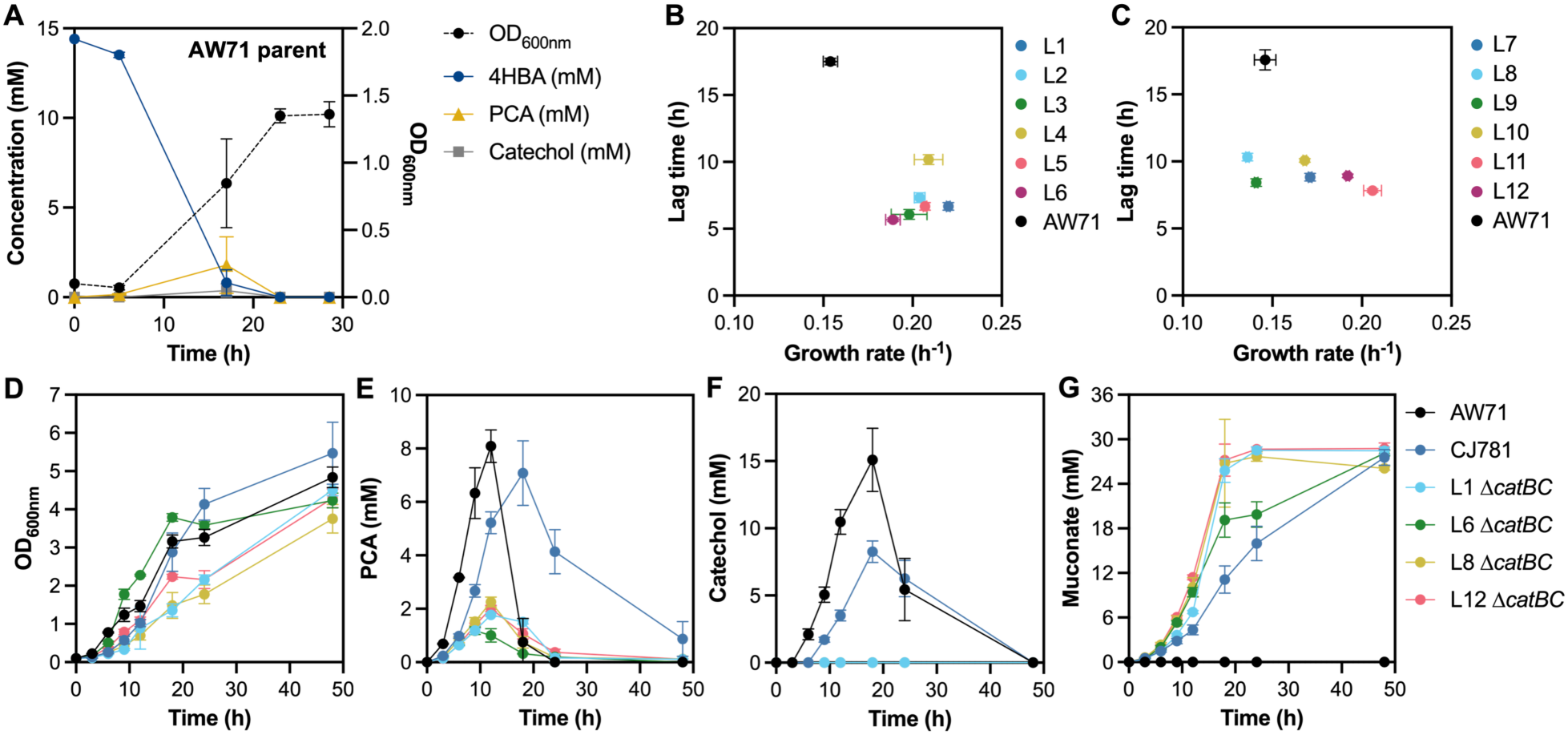
TALE alleviated bottlenecks at PCA and catechol, enabling improved muconate production from aromatic compounds. **(A)** The parent strain for evolution, AW71, accumulated PCA and catechol as intermediates when cultivated in M9 minimal medium with 15 mM 4HBA. **(B,C)** Evolved populations (lineage 1-6, L1-L6; lineage 7-12, L7-L12) exhibited improved growth rates and reduced lag times, relative to the parent strain, AW71; **(B)** shows data for populations evolved in M9 + 30 mM PCA, and **(C)** shows data for populations evolved in M9 + 30 mM 4HBA. The *catBC* genes were deleted in two evolved isolates per condition (L1 and L6 from TALE on PCA; L8 and L12 from TALE on 4HBA). When the resulting strains were cultivated in M9 + 30 mM 4HBA with glucose to support growth, they exhibited **(D)** changes in growth (OD_600nm_), reductions in **(E)** PCA and **(F)** catechol accumulation, and **(G)** improved muconate production from 4HBA, relative to AW71 and CJ781. In all plots, error bars indicate the standard deviation from the mean of three biological replicates. Numerical data for growth rate and lig time calculations are available in **Supplementary File 1**.

To assess whether the growth improvements obtained through ALE affected the PCA catabolic pathway in a manner that also affected muconate production, *catBC* was re-deleted in four evolved isolates (L1 and L6, evolved on PCA, and L8 and L12, evolved on 4HBA). Each Δ*catBC* isolate exhibited more rapid muconate production, relative to CJ781, in shake flask cultures supplemented with 4HBA (**Fig. 2D-G**). The Δ*catBC* isolates also showed much lower accumulation of PCA and no detectable accumulation of catechol in supernatants, indicating successful debottlenecking of these metabolic steps. A similar, but less substantial, improvement was observed for the Δ*catBC* isolates in cultures supplemented with 30 mM PCA (**Fig. S5A**). When evolved isolates were cultivated with *p*-coumarate, the catechol bottleneck was eliminated and a small reduction of the PCA bottleneck was observed, but no improvement in muconate production was realized, potentially because the additional steps required to catabolize *p*-coumarate (Jiménez et al., 2002) slowed the flow of carbon through these bottlenecks (**Fig. S5B**). Together, these results demonstrate that evolution selected for mutants whose growth was no longer impaired by accumulation of metabolic intermediates, and that improvements in AroY-dependent growth translated to faster muconate production upon deletion of *catBC*.

### 2.2. Mutations led to transcriptional changes in evolved strains

To investigate the genetic determinants of evolved phenotypes, whole-genome sequencing was conducted on AW71 and each of the evolved populations and isolates. A standardized analysis pipeline (see Methods) identified mutations in each sample, relative to the AW71 parent (**Supplementary File 1**; **Supplementary File 2**). Mutations occurred with remarkably high frequency in the *catA1* locus (PP_3713), with every isolate from the final passage of each of the twelve lineages exhibiting at least one mutation in this region (**Fig. 3A**). In addition to diverse single-nucleotide polymorphisms (SNPs) elsewhere in the genome, some lineages exhibited amplifications (2-5 fold) of large regions ranging from 11-300 kb in length. In one case (the final isolate from L9), a ∼300 kbp amplification spanned the *catA1* gene, further emphasizing the strong selection for mutation of this gene. Mutations to the gene encoding the surface-adhesion protein, LapA (PP_0168) (Espinosa-Urgel et al., 2000), and amplifications spanning the PP_3820-PP_3869 prophage region (Martínez-García et al., 2014) were also observed across both substrates. Isolates from lineages evolved on PCA exhibited a greater number of total mutations than those evolved on 4HBA (**Fig. S6**). Several convergent mutations occurred in intergenic regions, including SNPs in potential promoter regions for *vanK* (PP_3740), which encodes a PCA transporter (Wada et al., 2021), and PP_2553, which encodes a putative MFS transporter.

**Figure 3.**
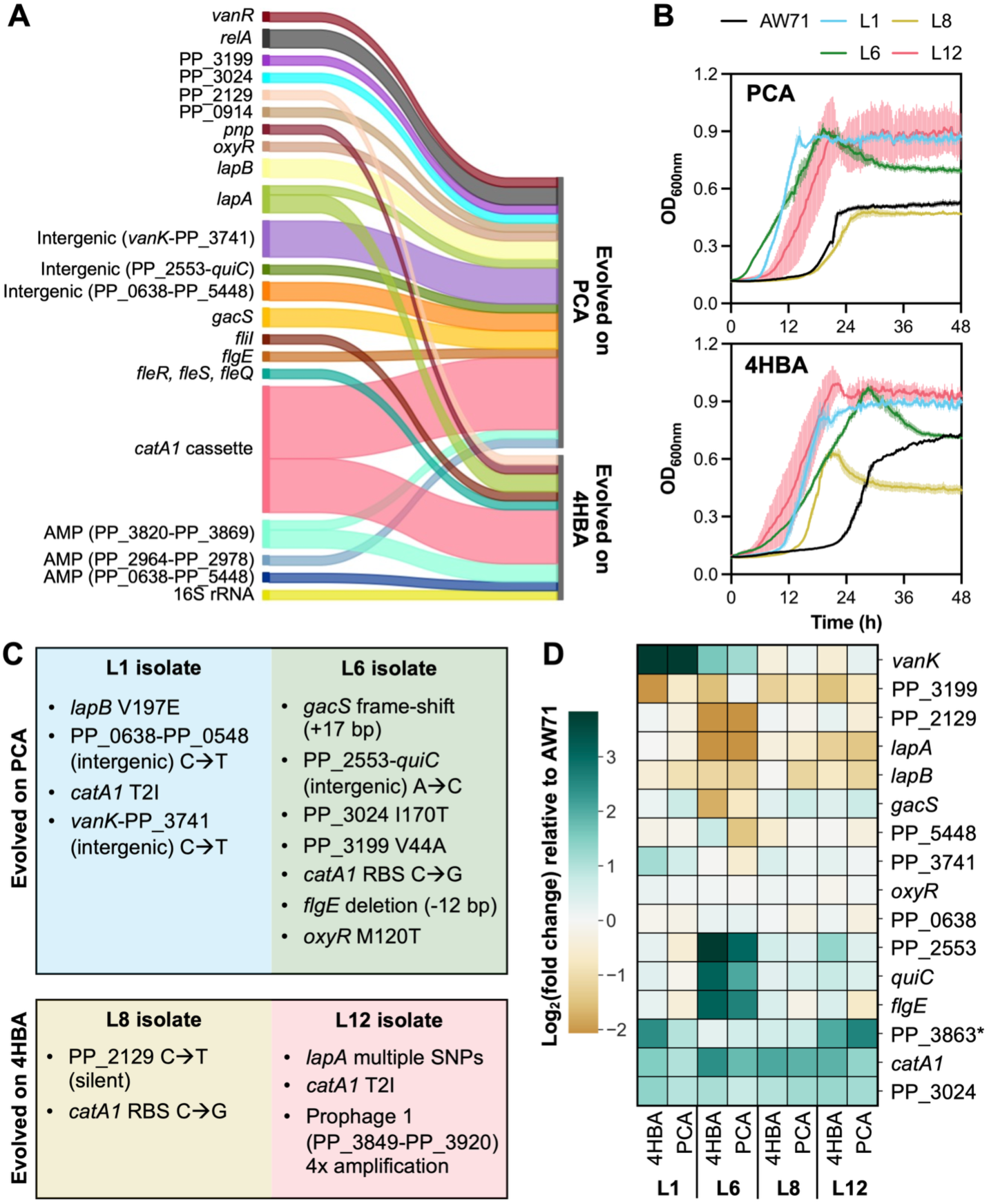
Mutation characteristics of evolved AW71 isolates. **(A)** Convergent mutations for all evolved isolates (L1-L12; **Supplementary File 2**). Line thickness corresponds to the number of times a mutation was observed. “Intergenic” indicates that a SNP was observed between the two indicated genes, and “AMP” refers to amplification of the indicated region. **(B)** The growth of evolved isolates (L1 and L6, evolved on PCA; L8 and L12, evolved on 4HBA) was compared to the AW71 parent in microtiter plates using M9 minimal media with 30 mM PCA or 30 mM 4HBA as the sole source of carbon and energy. Shading indicates the standard deviation from the mean of three biological replicates. **(C)** Mutations observed in each of the four isolates selected for transcriptomics analysis. **(D)** Transcript abundance for selected genes in each evolved isolate relative to AW71 (**Supplementary File 3**). Strains were grown to mid-exponential phase in shake flasks with M9 minimal medium and 30 mM 4HBA or 30 mM PCA prior to cell collection, library preparation, and RNAseq (see Methods). *PP_3863 is shown as a representative gene from the amplified region spanning PP_3849-PP_3920 in L12.

To elucidate the effects of ALE-derived mutations on gene expression, global transcriptomic analysis was performed on each of four isolates (L1 and L6, evolved on PCA, and L8 and L12, evolved on 4HBA), relative to the AW71 parent. These isolates generally exhibited improved growth on both substrates (**Fig. 3B**), but they displayed diverse mutations (**Fig. 3C**). Cells were collected at mid-exponential phase from cultivations in minimal medium with 30 mM PCA or 30 mM 4HBA, and RNA sequencing and relative transcript abundance quantitation were performed according to a previously described workflow (Holmes et al., 2024).

Mutations in each isolate corresponded to concomitant changes in gene expression (**Fig. 3D**). For example, isolate L1 contained a SNP in the promoter region for *vanK*, and this gene exhibited a >27-fold increase in expression in L1 relative to AW71 during growth with both substrates (**Supplementary File 3**). Isolate L6 contained a SNP in the intergenic region between PP_2553 and *quiC* (PP_2554), and we observed elevated expression of both genes. As expected from the strong selection for mutations in *catA1*, transcription of this gene was significantly elevated in each of the four isolates and on both substrates. Apart from expression changes linked to mutations, all evolved isolates exhibited significantly increased expression of the PP_2034-PP_2037 transcriptional unit and significantly decreased expression of the PP_3425-PP_3427 transcriptional unit (**Supplementary File 3**; **Fig. S7**). These operons contain *benE-I* (PP_2035), annotated as a benzoate transporter gene, and *mexEF*:*oprN*, which encodes an RND efflux pump, suggesting that metabolite transport may be modulated in the evolved isolates, relative to AW71.

### 2.3. Mutations to the *catA1* cassette alleviated metabolic bottlenecks

The most common convergent mutations from ALE occurred in the *catA1* cassette (**Fig. 3A**). Each of the twelve evolved isolates in the experiment contained at least one mutation to this cassette (**Supplementary File 2**), and mutations took several forms (**Fig. 4A,B**). Four isolates contained a T2I mutation in the *catA1* CDS and seven isolates contained a mutation in the ribosome binding site (RBS) for *catA1*. Two additional clones exhibited a G27A mutation in the *catA1* CDS, but it was always observed in conjunction with the C◊G SNP in the RBS.

**Figure 4.**
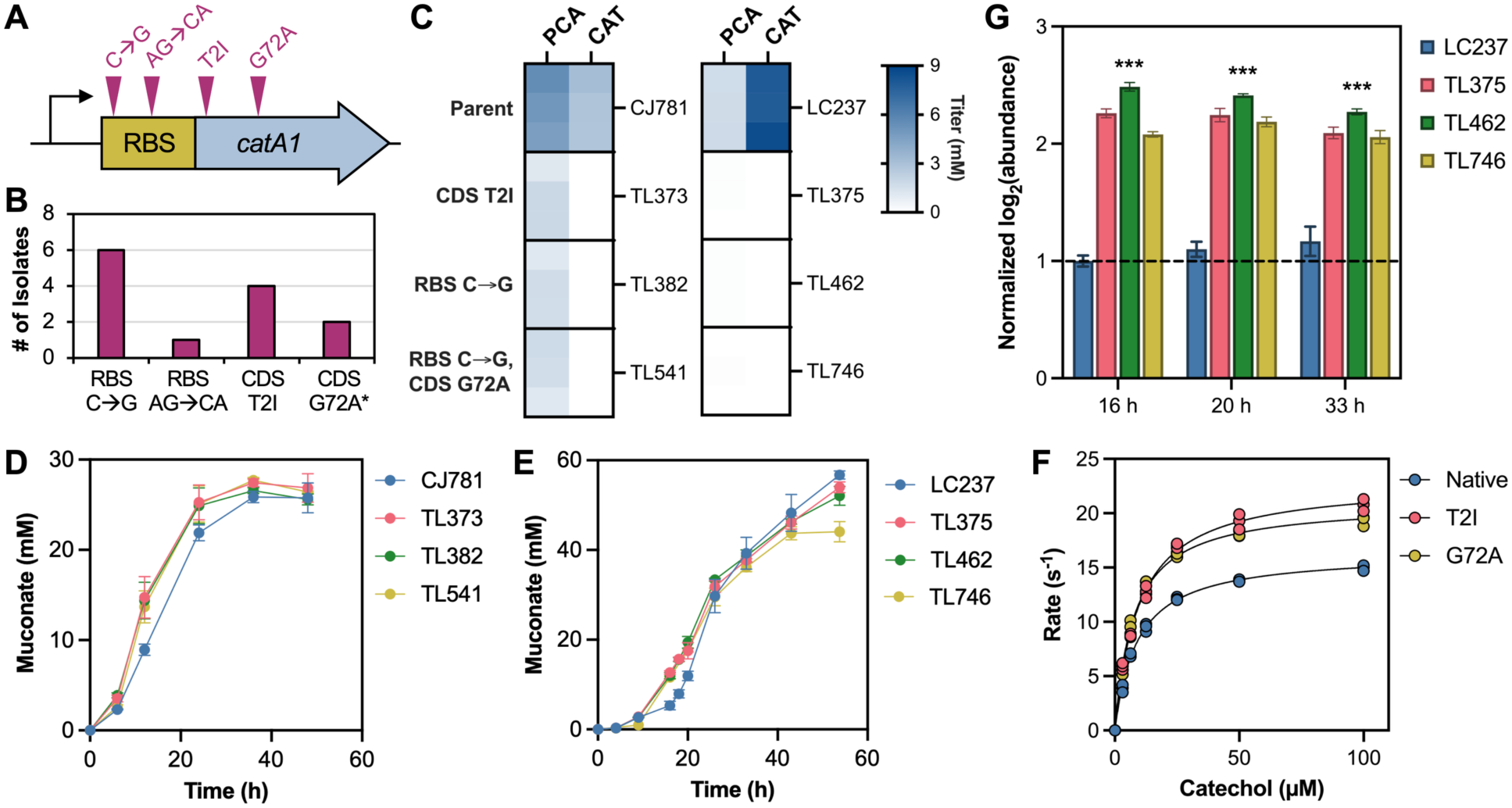
Analysis of mutations to the *catA1* cassette. **(A)** ALE-derived mutations to the *catA1* cassette were observed in the RBS and the CDS. **(B)** Occurrence count for each *catA1* mutation among all evolved isolates (L1-L12). *The G72A mutation was only observed in conjunction with the C◊G mutation in the RBS. **(C)** The three most common mutations in the *catA1* cassette were reverse-engineered into *P. putida* CJ781 (*left*) and *P. putida* LC237 (*right*). Strains were cultivated in triplicate in minimal media with 15 mM glucose + 30 mM 4HBA for CJ781-derived strains, or with mock hydrolysate sugars for LC237-derived strains. Heat maps indicate the maximum PCA and catechol (CAT) titer observed in each strain (at 12 h for CJ781-derived strains; at 20 h for LC237-derived strains). **(D,E)** ALE-derived mutations to the *catA1* cassette enabled faster muconate production rates, regardless of the strain background. Error bars indicate the standard deviation from the mean of three biological replicates. **(F)** Enzyme kinetics (**Table 2**; **Supplementary File 1**) were elucidated by measuring the rate of oxygen consumption by CatA1 and each of the mutants derived from ALE. Rates were corrected for the metal occupancy (Fe^3+^) of each enzyme. **(G)** Quantitative proteomics determined the abundance of CatA1 protein in strains LC237, TL375, TL462, and TL746 at each of the indicated timepoints during cultivation in shake flasks with minimal medium and mock hydrolysate sugars. All values were normalized to the abundance of CatA1 in LC237 at 16 h (dashed line). Error bars indicate the standard deviation from the mean of three biological replicates. \*\*\**p* < 0.001 for each of the three engineered strains, relative to LC237, in a two-tailed student’s t-test.

**Table 2.** Performance characteristics for the CatA enzymes described in this study. CatA activity was monitored via oxygen consumption at 25°C in air-saturated 25 mM HEPES, 50 mM NaCl, pH 7.5. Quantitative data for kinetic parameter calculations are included in **Supplementary File 1**.

| Enzyme | T<sub>m</sub> (°C) | Metal occupancy | *k*<sub>cat</sub><sup>app</sup> * (s<sup>-1</sup>) | *K*<sub>M</sub><sup>app</sup> (μM) | *k*<sub>cat</sub><sup>app</sup> / *K*<sub>M</sub><sup>app</sup> (s<sup>-1</sup> μM<sup>-1</sup>) | % relative to native |
| --- | --- | --- | --- | --- | --- | --- |
| CatA1 (native) | 51.1 ± 0.0 | 0.67 ± 0.06 | 16.3 ± 0.2 | 9.0 ± 0.3 | 1.8 ± 0.1 | 100 ± 6 |
| CatA1 T2I | 50.8 ± 0.3 | 0.47 ± 0.02 | 23.2 ± 0.3 | 9.6 ± 0.4 | 2.4 ± 0.1 | 133 ± 6 |
| CatA1 G72A | 52.2 ± 0.2 | 0.65 ± 0.04 | 21.1 ± 0.3 | 7.7 ± 0.4 | 2.7 ± 0.1 | 150 ± 6 |
| CatA2 (native) | 47.0 ± 0.5 | 0.24 ± 0.02 | 20.6 ± 0.3 | 7.9 ± 0.5 | 2.6 ± 0.2 | 150 ± 10 |

\* $k_{cat}^{app}$ is corrected for metal occupancy ( $Fe^{3+}$ )

To interrogate whether these mutations led to a conserved phenotype across strain backgrounds, we examined their effect on muconate production and intermediate metabolite accumulation from both lignin-derived aromatic and cellulosic sugar substrates. Specifically, we evaluated mutations in *P. putida* strain LC237 (Kim et al., 2026), which was engineered for muconate production from the cellulosic sugars (glucose, xylose, and arabinose), and *P. putida* strain CJ781 (Kuatsjah et al., 2022), which produces muconate from 4HBA when provided glucose as a carbon and energy source (**Table 1**). Strains AW71, CJ781, and LC237 share the terminal enzymatic steps for conversion of PCA to muconate (**Fig. 1A**), and they all accumulate PCA and catechol as metabolic intermediates (**Fig. 1B**, **Fig. 2A**, **Fig. S1**). We reverse-engineered convergent *catA1* mutations from ALE into the CJ781 and LC237 background strains, respectively generating strains TL373 and TL375 (T2I mutation to *catA1* CDS), strains TL382 and TL462 (C◊G SNP in *catA1* RBS), and strains TL541 and TL746 (C◊G SNP in *catA1* RBS plus G72A mutation to *catA1* CDS) (**Table 1**). When cultivated on relevant substrates, all engineered strains exhibited reductions in PCA and catechol accumulation (**Fig. 4C**, **Figs. S8-S9**) and increases in the muconate production rate (**Fig. 4D,E**). The combination of all three mutations was also evaluated in the CJ781 background (MG007; **Table 1**), but it did not exhibit significant improvements in intermediate accumulation or muconate production, relative to the individually reverse-engineered strains (**Fig. S8**). Together with the transcriptomics data, these results suggest that increased expression of *catA1* improves metabolic flux through PCA and catechol toward muconate, regardless of the growth substrate and strain background.

Mutations in *catA1* increased its expression in the evolved isolates, but it remained unclear whether improved muconate production and reduced intermediate accumulation stemmed from an increase in CatA protein stability, activity, or abundance. We thus analyzed the melting temperature (T_m_) for each of the mutants derived from ALE (CatA1 T2I and CatA1 G72A), relative to that of native CatA1 and native CatA2, but only minor T_m_ differences were observed among the variants (**Table 2**, **Fig. S10**), suggesting that protein stability is not the major driver of the improvements, at least as inferred from T_m_ alone. We also performed steady-state kinetics assays on each of the CatA1 mutants derived from ALE, as well as native CatA1 and native CatA2 (**Fig. 4F**, **Table 2**). Each of the mutants showed a modest increase in catalytic efficiency relative to the native CatA1 enzyme, with most of the efficiency gains originating from increased apparent turnover number (k_cat_^app^).

We subsequently conducted further experiments to evaluate the role of *catA1* mutations exclusively in the *P. putida* LC237 background, since it displayed higher accumulation of PCA and catechol intermediates than CJ781 under the conditions tested (**Figs. S8-S9**). First, we conducted quantitative proteomic analysis of CatA1 and CatA2 in strain LC237 during growth with mock hydrolysate sugars (93 mM glucose, 47 mM xylose, and 8 mM arabinose, representative of the ratio found in corn stover hydrolysate (Chen et al., 2016)). The *P. putida* genome encodes two catechol 1,2-dioxygenase enzymes, and CatA2 has been proposed as a “metabolic safety valve” in addition to CatA1 (Jiménez et al., 2014). As expected, the level of CatA2 did not change substantially between LC237 and the engineered strains, and its overall abundance was much lower than that of CatA1 (**Fig. S11**). However, each of the strains harboring mutations derived from ALE exhibited >6-fold higher levels of CatA1, relative to those in LC237 (**Fig. 4G**; **Supplementary File 4**). *In vivo*, the substantial increase in CatA1 protein levels in mutant strains likely overshadowed the relatively small activity differences observed in the CatA1 enzyme variants, but kinetic improvement of the mutant enzymes may also contribute synergistically to improved phenotypes.

### 2.4. Reverse engineering of other mutations contributed minor changes to phenotype

Evolved strains of AW71 developed an array of mutations but, to our surprise, none were directly observed in the *aroY* cassette. Parallel work with aromatic substrates suggested that increased expression of *aroY* alone is insufficient to overcome the metabolic bottlenecks in the pathway (Wilkes et al., 2026), so we examined this effect with sugar substrates in LC237. The synthetic *aroY:ecdB* operon was modified to increase the strength of the *aroY* RBS in LC237, resulting in strain ACB441 (**Table 1**). ACB441 contained a higher level of AroY protein than LC237, as measured by quantitative proteomics (**Fig. S12A**). PCA accumulation decreased in ACB441 compared to LC237, but catechol remained a dowstream bottleneck and muconate levels were unchanged (**Fig. S12B,C**). This observation suggested that increased expression of the downstream enzyme, CatA, contributed more strongly to the debottlenecking effect than AroY, and hence why our ALE campaigns did not select for mutations to AroY.

Although mutations to the *catA1* cassette were observed most frequently, ALE also resulted in a diverse range of non-intuitive mutations across the AW71 genome (**Fig. 3A**). We sought to examine the growth effect of a SNP between PP_2553 and *quiC* (PP_2554, a 3-dehydroshikimate dehydratase; **Fig. 1A**), which resulted in elevated expression of both genes in evolved strains (**Fig. 3D**, **Supplementary File 2**). We also noted some sequence ambiguity (identity <60% at a single nucleotide position) in the *dxs* (PP_0527) and *pdxJ* (PP_1436) genes in samples from the L8 lineage. The *dxs* gene encodes 1-deoxyxylulose-5-phosphate synthase, the first committed step of the MEP pathway for biosynthesis of dimethylallyl-diphosphate, an essential component of the PrFMN cofactor for AroY (Hernandez-Arranz et al., 2019; Wang et al., 2018). The pyridoxine 5’-phosphate synthase encoded by *pdxJ* acts on the product of Dxs, presenting a potential adjustment point for metabolic flux through these pathways (**Fig. 5A**). To investigate the effect of these mutations in a muconate-producing strain, as well as their generalizability to strains beyond those that utilize aromatics, we constructed a strain overexpressing the *dxs:dxr* operon under control of the strong P_tac_ promoter (strain ACB429; **Table 1**), and a strain harboring the intergenic PP_2553-*quiC* mutation (strain TLH070; **Table 1**). The resulting strains were cultivated in minimal media with mock hydrolysate sugars. Despite their predicted benefits to the pathway, neither the *dxs:dxr* overexpression strain nor the intergenic PP_2553-*quiC* mutant resulted in a substantial benefit for growth, muconate production, or PCA reduction (**Fig. 5B-D**, **Fig. S13**).

**Figure 5.**
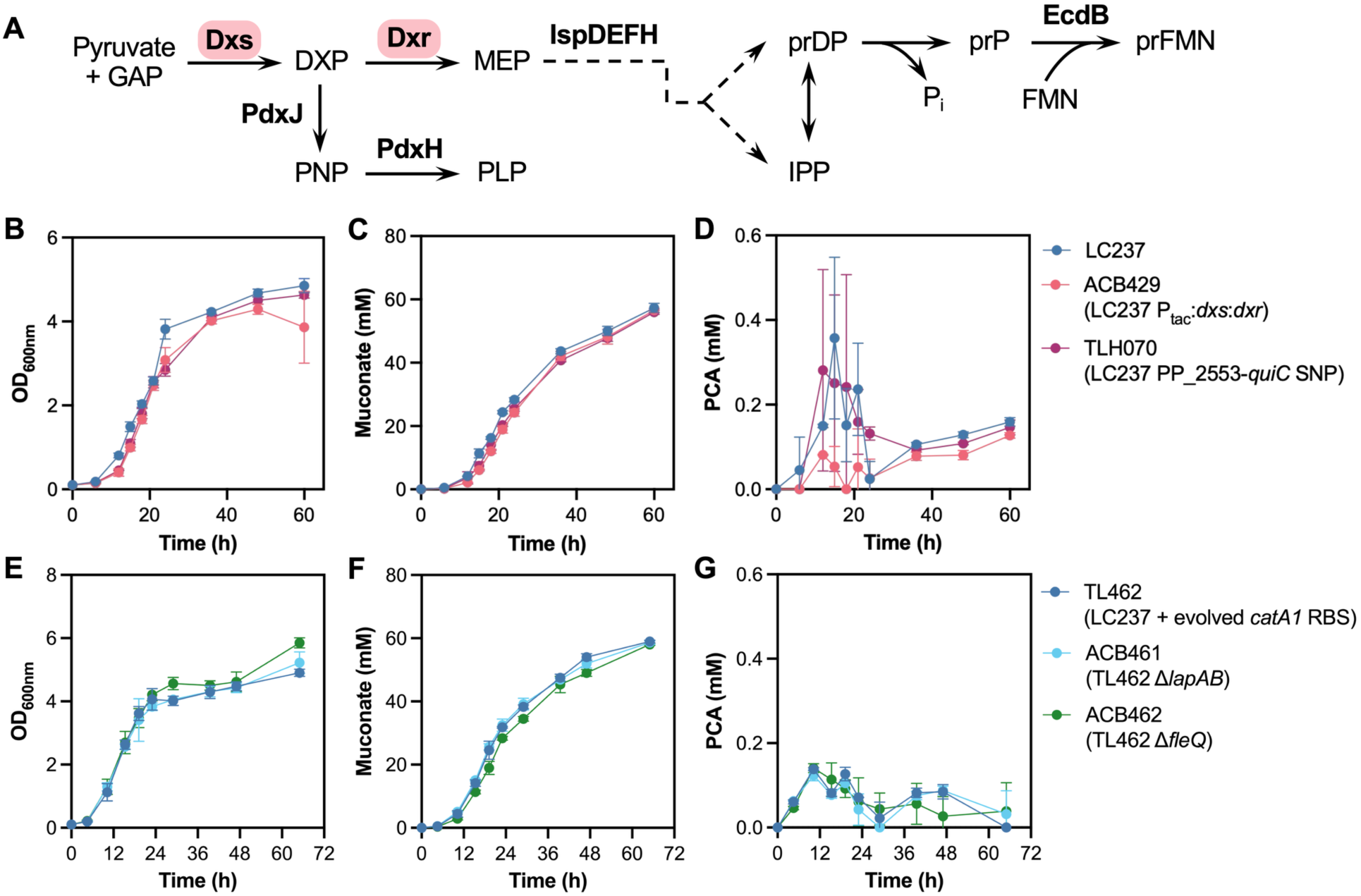
Evaluation of other ALE-derived mutations in *P. putida* LC237. **(A)** Generation of the AroY cofactor, prFMN. Enzymes in colored ovals were modified in strain ACB429. Variants of LC237 and TL462 harboring mutations derived from ALE were cultivated in shake flasks with minimal medium containing 25 g/L mock hydrolysate, with measurement of **(B,E)** cell density (OD_600nm_), **(C,F)** muconate titer, and **(D,G)** PCA accumulation. Error bars indicate the standard deviation from the mean of three biological replicates. Abbreviations: GAP, glyceraldehyde 3-phosphate; DXP, 1-deoxy-D-xylulose 5-phosphate; MEP, 2-C-methyl-D-erythritol 4-phosphate; prPP, prenyl diphosphate; IPP, isopentenyl diphosphate; prP, prenyl phosphate; FMN, flavin mononucleotide; PNP, pyridoxine 5’-phosphate; PLP, pyridoxal 5’-phosphate.

Other changes from ALE were somewhat intuitive; for example, the loss-of-function mutations observed in the *lapAB* cell-surface adhesion complex and its regulators, *gacS* and *fleQ*, aligned with previously observed fitness benefits for deletion of these genes in *P. putida* (Bentley et al., 2020; Borchert et al., 2023; Eng et al., 2021; Martínez-Gil et al., 2014; Werner et al., 2025). Strain TL462, the variant of LC237 containing a C◊G SNP in the RBS for *catA1* (**Table 1**), was further modified to add deletions either in *lapAB* (ACB461) or *fleQ* (ACB462), but this combination failed to show a substantial change in growth, muconate production, PCA accumulation, or substrate consumption, relative to TL462 (**Fig. 5D-F**, **Fig. S14**). Similar results were obtained for deletion of *lapAB* or *fleQ* alone in the original LC237 background (strains ACB458 and ACB459; **Fig. S15**). Overall, these investigations affirmed the strong selection for mutations that increased the abundance of catechol 1,2-dioxygenase, and showed that no other gene modification examined here induced a phenotypic change as effectively as those observed with *catA1*.

## 3. Discussion

Heterologous expression of *aroY* enables strains of *P. putida* to convert PCA to catechol (Johnson et al., 2016; Vardon et al., 2015). In this work, we used ALE to improve the efficiency of the AroY route to muconate, and evolved strains exhibited significant increases in growth while eliminating detectable metabolic bottlenecks at PCA and catechol. Reductive decarboxylation of PCA to catechol serves as a critical component of native aromatic catabolism in a range of microorganisms, including bacteria such as *Klebsiella* and *Enterobacter* as well as fungal species such as *Arxula adeninivorans* (Grant and Patel, 1969; Matsui et al., 2006; Meier et al., 2017). Bacterial species, including those engineered in this study, typically utilize the UbiD-type PCA decarboxylase system wherein a flavin prenyltransferase generates the prFMN cofactor required for efficient activity of the decarboxylase (Payne et al., 2015; White et al., 2015). We originally hypothesized that PCA accumulation in engineered strains of *P. putida* arose from poor AroY activity and/or high demand for the prFMN cofactor in the context of *aroY* overexpression, but ALE overwhelmingly selected for mutations that increase the transcription, translation, and turnover of the next enzyme in the pathway, CatA, even though *catA1* was already under control of the strong P_tac_ promoter in the initial ALE strain, AW71 (**Table 1**). This suggested that CatA activity represented the rate-limiting step in muconate production from PCA in these strains. Indeed, modification of *P. putida* strains to increase the activity of CatA1—through elevated expression and/or catalytic enhancement—fully eliminated the bottlenecks at both PCA and catechol, increasing muconate production in multiple strains that rely on the AroY decarboxylase, including those that produce muconate from lignin-related aromatic compounds and those that produce muconate from cellulosic sugars. As an alternative strategy to the ALE approach taken here, AroY could potentially be exchanged for cofactor-less eukaryotic decarboxylases such as AGDC1 from *A. adeninivorans* (Brückner et al., 2018); however, our attempts to express these enzymes in *P. putida* failed to produce active strains, and the data in this study indicate that CatA is the more critical enzyme for pathway optimization, with PCA accumulation likely occurring as a consequence of equilibrium with catechol accumulated downstream. In parallel work utilizing multi-omics to improve muconate production from lignin-related aromatics during glucose co-feeding, we arrived at a similar conclusion: increased expression of CatA2 and, to a lesser extent, AroY enabled increased muconate titers, rates, and yields (Wilkes et al., 2026). Related work in our group has also leveraged elevated expression of CatA1 as a strategy for improving muconate titers, rates, and yields from biomass hydrolysate sugars in bioreactors (Kim et al., 2026).

This study illustrates the utility of ALE in pinpointing critical mutations for improved growth and highlights the use of replicates to enhance confidence in the relative impact of each mutation. Even though ALE selected for several distinct modifications to the *catA1* cassette, these resulted in similarly elevated levels of CatA1 protein, reflective of intrinsic tradeoffs between growth rate and protein translation rate in bacteria (Maitra and Dill, 2014; Weiße et al., 2015). In practice, each unique *catA1* mutation from ALE may have driven the abundance of this enzyme to the same optimum level, explaining the relatively consistent CatA1 protein level observed across reverse-engineered strains. Evolution also selected for mutations that modestly accelerated the catalytic efficiency of CatA1, but these gains were likely eclipsed by large increases in protein abundance. Nevertheless, the increased catalytic efficiency of CatA1 variants (T2I and G72A) could be leveraged to decrease the expression of *catA1* for conservation of cellular resources, or they could serve as superior biocatalysts in separate applications, such as cell-free systems that require high rates for effective product generation.

The success of this ALE campaign illustrates the importance of strain choice and experimental design in selecting for critical mutations. Previously, we attempted evolution of an AroY-dependent *P. putida* strain on *p-*coumarate, but growth with this substrate did not result in considerable accumulation of PCA or catechol as intermediates. TALE of AW71 with PCA and 4HBA imposed selective pressure directly at the metabolic bottleneck, leading to strong selection for mutations which alleviated accumulation of intermediates. Still, some mutations were not obviously related to the AroY pathway and likely arose from use of the aromatic substrates in ALE; e.g., mutations to *vanK* indicated that this transporter plays a role in uptake of PCA and 4HBA in addition to vanillate (Wada et al., 2021). In other cases, reverse engineering of single mutations failed to elicit an increase in growth or muconate production (**Fig. 6**). This suggested a role for epistasis, in which modification of two or more sites is required to confer a phenotypic effect (Johnson et al., 2023).

## 4. Conclusions

In summary, this study leveraged the selective power of ALE to identify *catA1* as a critical gene target for optimized growth in strains of *P. putida* engineered for muconate production. We demonstrated the importance of strong *catA1* expression for metabolic debottlenecking in multiple strain backgrounds, and future work will determine whether the beneficial effects of the mutations described here can be translated to additional strain architectures in *P. putida* and beyond. Together, these efforts accelerate the development of microbial platforms for production of a key commodity chemical from sustainable feedstocks, laying the groundwork for bioprocess development and scale-up.

## 5. Methods

### 5.1 Strain construction and cultivation

#### 5.1.1. Strain and plasmid construction

The bacterial strains constructed in this study are listed in **Table 1**, with construction details described in **Table S1**. Plasmids are listed in **Table S2**, and oligonucleotides in **Table S3**. For some entries in **Table S2**, plasmids were synthesized by Twist Biosciences. For the remainder, plasmids were constructed in-house using fragment amplification by PCR with Q5® High-Fidelity 2X Master Mix (New England Biolabs, NEB). Following PCR, fragments were cleaned up with the NucleoSpin Gel and PCR Clean-Up kit (Macherey-Nagel) and underwent Gibson assembly with NEBuilder® HiFi DNA Assembly Master Mix (NEB). Assembly reactions were transformed into NEB® 5-alpha F’*I^q^ E. coli* (NEB) and plated on LB agar with required antibiotic (see **Table S2**). Plasmids were sequenced by Oxford Nanopore (Plasmidsaurus). For heterologous protein expression, plasmids were transformed into NEB® BL21(DE3) competent *E. coli* according to the manufacturer instructions. For genome editing of *P. putida* strains with pK18(m)sB-based plasmids, the parent strain was grown overnight in LB medium and made electrocompetent according to an established protocol (Choi et al., 2006). Cells were electroporated with >400 ng plasmid using a Bio-Rad GenePulser (1.6 kV, 25 µF, 200 Ω) and immediately transferred to SOC Outgrowth Medium (NEB). Outgrowth occurred at 225 rpm and 30°C for 60-120 min, and cultures were selected on LB agar with 50 mg/L kanamycin. Transformants were selected again on LB + 50 mg/L kanamycin and counter-selected twice on YT + 25% sucrose (10 g/L tryptone, 5 g/L yeast extract, 250 g/L sucrose, 18 g/L agar), according to a previously established protocol (Johnson and Beckham, 2015). Genomic edits were confirmed by PCR amplification of the region of interest with MyTaq^TM^ Red Mix (Meridian Bioscience) followed by Oxford Nanopore sequencing of the product (Plasmidsaurus).

#### 5.1.2 Media preparation

LB (Miller) served as the rich medium for strain maintenance and selection in this study. For growth experiments with defined carbon sources, M9 minimal medium was prepared with 0.1 mM CaCl_2_, 18 µM FeSO_4_, 2 mM MgSO_4_, 3 g/L KH_2_PO_4_, 0.5 g/L NaCl, 6.8 g/L Na_2_HPO_4_, 1 g/L NH_4_Cl, and the listed concentration(s) of carbon growth substrates. For ALE experiments, M9 medium was prepared at 2X concentration and mixed in a 1:1 ratio with the relevant aromatic substrate at 2X concentration. Aromatic compound stocks were prepared by dissolving PCA, 4HBA, or *p*-coumarate (all from Sigma-Aldrich) in water and then titrating the pH to neutral with 4 M NaOH. For experiments with mock hydrolysate sugars, M9 medium was prepared with 93 mM glucose, 47 mM xylose, and 8 mM arabinose – a typical sugar ratio observed in corn stover hydrolysates (Chen et al., 2016). For washing cells, 1X M9 salts contained 3 g/L KH_2_PO_4_, 0.5 g/L NaCl, 6.8 g/L Na_2_HPO_4_, and 1 g/L NH_4_Cl. All M9 media were sterilized through a 0.2 µm bottletop filter prior to use.

#### 5.1.3. Plate reader cultivations

Test tubes containing 5 mL LB medium were inoculated with strains of interest, directly from glycerol stocks. Tubes were incubated at 225 rpm and 30°C overnight. 1 mL of each culture was centrifuged at 2,500x*g* for 3 min and pellets were resuspended in 1 mL 1X M9 salts. Wells of Honeycomb2 microtiter plates (Growth Curves Ltd.) were filled with 200 µL M9 minimal media containing carbon source(s) of interest, and then triplicate wells were inoculated with 1 µL each of relevant culture suspension. Plates were incubated in a BioscreenC Pro (Growth Curves Ltd.) at 30°C with continuous shaking at maximum amplitude. Growth rates were calculated by linear regression during exponential phase, and lag time was defined as the growth time required to reach OD600nm = 0.2.

#### 5.1.4. Shake flask cultivations

Baffled flasks containing 25 mL LB medium were inoculated with strains of interest, directly from glycerol stocks. Flasks were incubated at 225 rpm and 30°C overnight. The next day, each overnight culture was diluted into 25 mL fresh LB medium to an OD_600nm_ of ∼0.2. Flasks were incubated at 225 rpm and 30°C for 4 h, and then precultures were centrifuged at 2,500x*g* for 6 min. Alternatively, for the CJ781-derived aromatic strains shown in **Fig. S8**, overnight precultures were processed directly without a second serial passage. Supernatants were decanted and pellets were resupended in 25 mL/each 1X M9 salts, and the centrifuge step was repeated. A second wash was performed, and then cell pellets were resuspended in 2.5 mL 1X M9 salts. The OD_600nm_ of each suspension was measured on a Gensys (Thermo Scientific) spectrophotometer, and these were used to inoculate baffled flasks containing 25 mL M9 minimal medium with carbon sources as indicated in each experiment. Flasks were incubated at 225 rpm and 30°C. At indicated time points, 800 µL of culture was withdrawn from each flask; 50 µL of cell suspension was utilized to measure OD_600nm_. The remaining cells were pelleted and supernatants passed through 0.2 µm filters into glass vials for metabolite quantification. The pH of each flask was also monitored at each sampling time using a miniature pH meter (HORIBA LAQUAtwin pH-33), and when necessary, sterile 1 M NaOH was used to adjust the culture to pH 7.

#### 5.1.5. Metabolite quantification

Muconic acid, aromatic compounds, and sugars were quantified from cell supernatants using ultra-high performance liquid chromatography (UHPLC) as described in published protocols.io methods (Alt et al., 2024; Woodworth et al., 2024).

### 5.2 Tolerance ALE

Prior to the start of TALE, strain AW71 was struck for isolation onto LB agar and incubated at 30°C until colonies formed. Each of six individual test tubes was filled with 5 mL LB medium, and each tube was inoculated with a single colony of AW71 (biological replicates). Tubes were incubated in an angled rack at 225 rpm and 30°C for 18 h. The following day, optical density at 600 nm (OD_600nm_) was measured for each overnight culture on a spectrophotometer (Thermo Scientific). Overnight cultures were used directly (no washing) to inoculate ALE cultures to an initial OD_600nm_ of 0.03. ALE cultivations were carried out in sterile test tubes with 5 mL M9 minimal medium containing 15 mM PCA or 15 mM 4HBA, incubated in an angled rack at 225 rpm and 30°C. The following day, OD_600nm_ was measured in the test tube using a Spectronic 601 tube spectrophotometer (Milton Roy). When OD_600nm_ values reached ∼1.0, 50 µL of each culture was passaged into fresh media in sterile test tubes. The timing and OD_600nm_ for each passage are recorded in **Table S4**. Evolving populations from each lineage were saved periodically throughout the TALE as glycerol stocks (20% w/v glycerol, -80°C) for later recovery. Every two weeks (see **Table S5**), the concentration of aromatic substrate was increased by 2.5 mM, and TALE continued until the final concentration of 30 mM was reached (44 passages in total). The final population from each lineage was stored as a glycerol stock, and 20 µL of liquid culture from each population was struck for isolation on M9 agar containing relevant aromatic compound (15 mM PCA or 15 mM 4HBA). Plates were incubated until single colonies formed, and then a single colony from each lineage was inoculated into an overnight culture in 5 mL LB medium. Each of the resulting isolates was also stored as a glycerol stock.

### 5.3. Whole genome sequencing and transcriptomics analysis

#### 5.3.1. Whole genome sequencing and mutation calling in AW71

Prior to the start of ALE, strain AW71 was inoculated directly from a glycerol stock into 5 mL LB medium and grown overnight at 225 rpm and 30°C. A 1 mL cell pellet was withdrawn, stabilized in DNA/RNA Shield Solution (Zymo Research), and sent for Illumina whole genome sequencing at ∼160X depth using 2x150 bp reads (Azenta Life Sciences). A reference genome for AW71 was constructed by manually modifying the *P. putida* KT2440 wild-type consensus genome to reflect each deletion and heterologous cassette. Read alignments were performed in Geneious software version 2023.0.4 using the Geneious mapper with medium sensitivity and five iterations. SNPs were called relative to the AW71 reference genome using a minimum coverage of 10 reads, minimum variant frequency of 65%, maximum variant P-value of 10^-6^, minimum strand-bias P-value of 10^-5^ when exceeding 65% bias, and bacterial default genetic code. The SNP table can be found in **Supplementary File 1**.

#### 5.3.2. Whole genome sequencing and mutation calling in evolved lineages

Evolved isolates and populations were inoculated directly from the glycerol stock into 5 mL LB medium and grown overnight at 225 rpm and 30°C. A 1 mL cell pellet was withdrawn, stabilized in DNA/RNA Shield Solution (Zymo Research), and sent for whole-genome sequencing by Oxford Nanopore (Plasmidsaurus). Reads were mapped to the AW71 reference genome using Geneious as described above, but with the medium-low sensitivity setting. For evolved populations, SNPs were called using a minimum variant frequency of 55%, while evolved isolates were called using a minimum variant frequency of 75%. Geneious was also used to identify regions of high coverage (>3 standard deviations from the mean; default settings) to predict genome amplification events. Mutation calls were then filtered to remove those that were also observed in the AW71 whole genome sequencing run, followed by a second round of manual filtering to ignore SNP calls caused by tandem repeat errors. The full table of mutations can be found in **Supplementary File 2**.

#### 5.3.3. Transcriptomics analysis

Baffled flasks containing 50 mL LB medium were inoculated with AW71 and four evolved isolates (L1, L6, L8, and L12), directly from glycerol stocks. Flasks were incubated at 225 rpm and 30°C for 18 h, and then cultures were centrifuged at 3500 xg for 10 min. Cell pellets were resuspended in 10 mL sterile 1X M9 salts and the OD_600nm_ of each was measured on a Gensys (Thermo Scientific) spectrophotometer. Each strain was used to inoculate three baffled flasks containing 25 mL M9 minimal medium + 30 mM PCA and three flasks containing M9 minimal medium + 30 mM 4HBA to initial OD_600nm_ of 0.1. Flasks were incubated at 225 rpm and 30°C, and OD_600nm_ was measured periodically. When cultures reached mid-exponential phase (OD_600nm_ ∼1.0-1.5, between 12 and 24 h), 2 mL of cells were pelleted at 6000 xg for 1 min, flash frozen on liquid nitrogen, and frozen at -80°C. Frozen pellets were sent to Azenta Life Sciences for RNA extraction, library preparation, and RNA sequencing (Illumina HiSeq, 2 x 150 bp reads, >25 M reads per sample). RNA sequencing data were analyzed using the KBase platform (https://www.kbase.us) (Arkin et al., 2018) according to a previously established approach (Holmes et al., 2024) utilizing Trimmomatic, Bowtie2, StringTie, and DESeq2 (Bolger et al., 2014; Langmead and Salzberg, 2012; Love et al., 2014; Pertea et al., 2015). Differential expression output files from DESeq2 are provided in **Supplementary File 2**.

### 5.4. Targeted proteomics

Proteins were extracted from the lyophilized cell pellets of engineered *P. putida* strains using the MPLEx method (Nakayasu et al., 2016). Proteins were trypsin-digested into peptides. Targeted proteomics was conducted utilizing established assays (peptides and transitions); comprehensive details on sample preparation and targeted proteomics assay development can be found in a previously published study (Gao et al., 2020). Crude heavy isotope-labeled internal standards were spiked in the peptide samples before liquid chromatography-selected reaction monitoring/mass spectrometry (LC-SRM/MS) analysis. Samples were analyzed using a Thermo Altis triple quadrupole mass spectrometer connected with a Neo Vanquish liquid chromatography system (Thermo Fisher Scientific). The LC utilized an in-house packed 100 μm i.d. × 20 cm column with ACQUITY UHPLC BEH 1.7 μm C18 particles and a reversed-phase chromatographic separation with mobile phases of 0.1% formic acid in water and acetonitrile. The data were acquired in scheduled SRM mode with a unit resolution of 0.7 FWHM for Q1 and Q3, and an optimized Q2 gas pressure of 1.5 mTorr. All LC-SRM data were imported into Skyline software (MacLean et al., 2010), and peak boundaries were manually reviewed to confirm accurate peak assignment and peak boundaries. The total peak area ratios between endogenous light peptides and their corresponding heavy isotope-labeled internal standards were exported from the Skyline software for subsequent analysis (**Fig. S16**). Data processing and quality control were performed at both peptide and protein levels using pmartR (Degnan et al., 2023). Data were log_2_-transformed, and outliers were identified and removed based on a Robust Mahalanobis Distance (RMD) procedure. Quantitative comparisons of log_2_-fold change were conducted via two-sample t-tests with Holm adjustments for multiple comparisons.

### 5.5 Protein expression and assays

#### 5.5.1. Protein expression and purification

Protein used for *in vitro* assays were recombinantly produced in *E. coli* BL21(DE3) (NEB) transformed with appropriate plasmids (**Table S2**). The cultures were grown in LB supplemented with 0.1 mg/mL ampicillin at 37°C at 200 rpm unless stated otherwise. A starter culture (40 mL media in 125 mL flask) was grown for 16 h and used to seed the main culture (1 L media in 2.5 L baffled flask) at 1% (v/v). The culture was induced with 1 mM isopropyl-β-D-thiogalactopyranoside and 0.5 mM ferric chloride upon reaching an OD_600nm_ of ∼0.7. The growth temperature was lowered to 18°C and the induced culture was grown for an additional 16 h. The resulting biomass was harvested by centrifugation (30 min at 4,000*xg*) and stored at -80°C until further processing.

Protein purification was performed using successive immobilized metal affinity chromatography and anion exchange chromatography steps. The frozen biomass was thawed and resuspended in ∼10 mL Buffer A (20 mM HEPES, 100 mM NaCl, pH 7.5) per liter of main culture used. Trace amounts (∼0.1 mg each) of lysozyme (Roche) and DNAseI (Roche) were added to cell suspensions to help with lysis and downstream processing. The cell suspension was lysed by sonication (Qsonica) and clarified by centrifugation (40 min at 16,000*xg*) followed by passaging through a 0.45 μm syringe-driven filter. The cleared lysate was applied to NiNTA resin (Goldbio). The resin was washed with Buffer A containing up to 40 mM imidazole. The dioxygenase was eluted with Buffer A containing 300 mM imidazole. The eluate was concentrated and buffer exchanged to imidazole-free Buffer A by successive dilution using centrifugal filtration (30 kDa, Millipore) for 20 min at 4,000*xg*. The concentrated protein was subsequently applied to a Source-Q resin column (Cytiva) and purification was carried out using an ÄKTA Pure fast-protein liquid chromatography system (Cytiva). The protein was eluted from the resin following a gradient of Buffer A containing up to 1 M NaCl. Fractions of interest, as identified from the spectroscopic profile and SDS-PAGE, were pooled, concentrated, and buffer exchanged to Buffer A as before. The resulting purified protein concentrate was flash frozen with liquid nitrogen and stored at -80°C until further use.

#### 5.5.2. Iron quantification assay

The iron content of purified CatA enzymes was analyzed using a modified protocol (Haigler and Gibson, 1990) adapted for microplate format following the formation of chromogenic complex between Ferene-S and Fe^2+^. Protein sample (80 μL) was denatured following the addition of HCl (12 N; 10 μL) and trichloroacetic acid (80%; 10 μL). The precipitated protein was removed by centrifugation. The supernatant was transferred to a fresh tube, diluted with sodium acetate (45% (w/v); 20 μL), and mixed with the Ferene-S solution (0.75 mM Ferene-S, 10 mM L-ascorbic acid, 45% (w/v) sodium acetate; 180 μL). The complex formation was monitored at 593 nm and compared to a standard curve of known ferric chloride concentrations. The metal occupancy was obtained by comparing the molar iron concentration from the assay above relative to the protein concentration determined using a bicinchoninic acid-based protein quantification kit (Pierce).

#### 5.5.3. Enzyme kinetics

Activity of each CatA enzyme was monitored by following the consumption of O_2_ using a Clark-type electrode, OXYG1+ oxygraph (Hansatech). The oxygraph was calibrated daily using air-saturated water and sodium hydrosulfite according to manufacturer instructions. The typical assay was conducted in air-saturated 25 mM HEPES, 50 mM NaCl, pH 7.5 at 25°C. Enzyme activity was monitored upon addition of purified protein (20 – 80 nM) to the buffer solution containing different concentrations of catechol (0-0.1 mM; final DMSO concentration < 0.5 % (v/v)). The reading was corrected with the background oxygen concentration prior to the addition of the enzyme. The apparent steady-state kinetic parameters were evaluated by fitting the Michaeles-Menten equation to the data.

#### 5.5.4. Differential scanning fluorimetry

The melting temperature of each CatA enzyme was determined using a previously described differential scanning fluorimetry method which follows the formation of fluorogenic signal upon protein denaturation (Norton-Baker et al., 2024). In short, the protein solution was mixed with Sypro Orange Dye (ThermoFisher) and subjected to a ramp cycle in a CFX96 Real-Time System (BioRad). The resulting fluorescence formation as a function of temperature was plotted and its derivative was used to determine the melting temperature. These measurements were performed in 20 mM HEPES, 100 mM NaCl, pH 7.5.

## CRediT authorship contribution statement

Conceptualization: A.C.B., C.W.J., and G.T.B. Experimentation: A.C.B., T.L.H., T.M.L., E.K., Y.G., M.A.G., Z.A.K., and A.Z.W. Visualization: A.C.B. Funding acquisition: C.W.J. and G.T.B. Supervision: A.C.B., A.Z.W., Y.M.K., C.W.J., and G.T.B. Writing—original draft: A.C.B. Writing—review and editing: A.C.B., T.L.H., T.M.L., E.K., A.Z.W., Y.M.K., C.W.J., and G.T.B.

## Acknowledgements

This work was authored by the National Laboratory of the Rockies for the U.S. Department of Energy (DOE) under contract no. DE-AC36-08GO28308. Funding was provided to M.A.G., A.Z.W., and G.T.B. by the U.S. DOE, Office of Critical Minerals and Energy Innovation, Alternative Fuel and Feedstocks Office (AFFO). A.C.B., T.L.H., T.M.L., Y.G., Y.M.K., C.W.J., and G.T.B. were additionally supported by AFFO for the Agile BioFoundry via contract no. DE-AC36-08GO28308. This work was also supported for A.C.B., E.K., Z.A.K., A.Z.W., and G.T.B. by the Center for Bioenergy Innovation (CBI), U.S. DOE, Office of Science, Biological and Environmental Research Program under Award Number ERKP886. The DOE Systems Biology Knowledgebase (KBase) is funded by the U.S. DOE, Office of Science, Office of Biological and Environmental Research. A portion of this work was performed at Pacific Northwest National Laboratory using EMSL, a DOE Office of Science User Facility sponsored by the Office of Biological and Environmental Research. The views expressed in the article do not necessarily represent the views of the DOE or the U.S. government. We thank Alex Benson and Kelsey J. Ramirez for their efforts in metabolite quantification and Adam M. Feist and Caroline R. Amendola for helpful discussions.

## Data Availability

Whole-genome and RNA sequencing read files from this study are available in the NCBI repository under Sequence Read Archive (SRA) BioProject ID number PRJNA1483726. Datasets supporting the results of this article are included in this article and its supplementary files. Additional data will be made available upon request.

