## Supplemental Information for "Overcoming protocatechuate and catechol accumulation in muconic acid production via adaptive laboratory evolution and metabolic engineering in *Pseudomonas putida*"

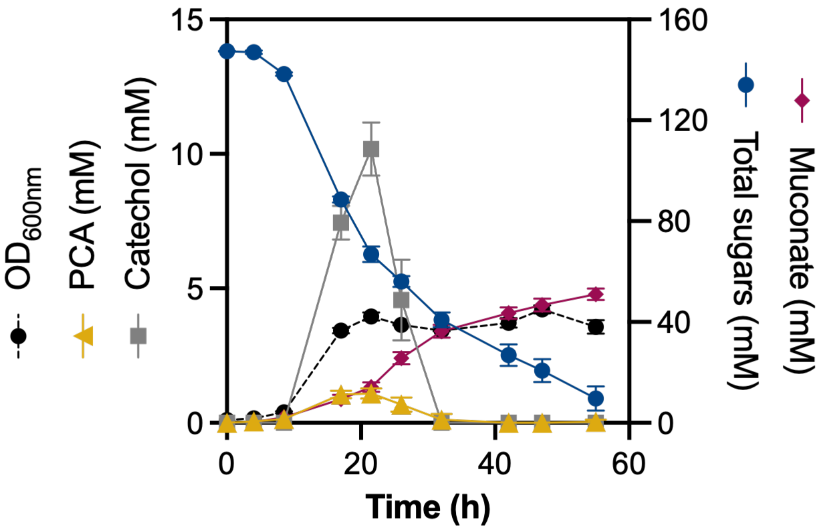

**Figure S1. *P. putida* LC237 accumulates PCA and catechol as metabolic intermediates.** The strain was cultivated in shake flasks in M9 minimal medium with 25 g/L hydrolysate sugars (93 mM glucose, 47 mM xylose, and 8 mM arabinose, where ratios were chosen to represent the sugars composition of corn stover hydrolysate). Error bars indicate the standard deviation from the mean of three biological replicates.

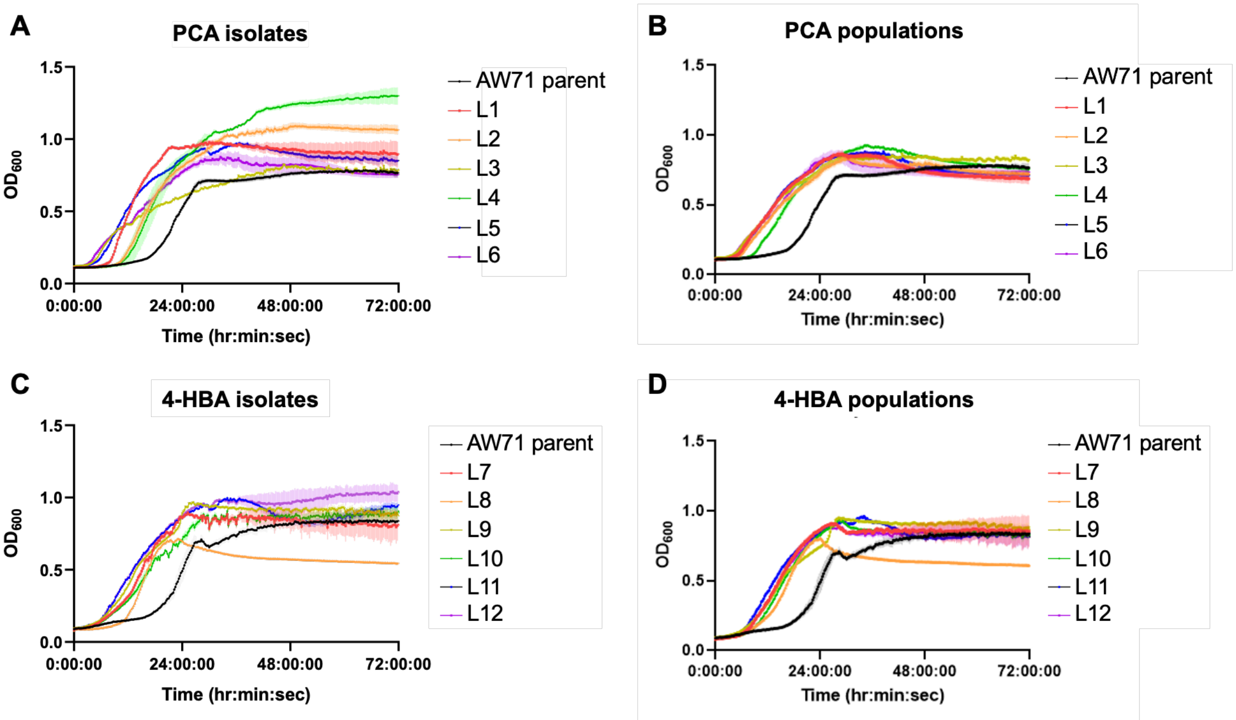

**Figure S2. Growth phenotypes of evolved strains from ALE. (A)** Clonally unique isolates were collected from each lineage of the TALE on PCA, and their growth was compared to that of the parent strain, AW71, in M9 minimal medium with 15 mM PCA. **(B)** Clonally unique isolates were collected from each lineage of the TALE on 4HBA, and their growth was compared to that of the parent strain, AW71, in M9 minimal medium with 15 mM 4HBA. **(C)** Populations samples were collected from each lineage of the TALE on PCA, and their growth was compared to that of the parent strain, AW71, in M9 minimal medium with 15 mM PCA. **(D)** Populations samples were collected from each lineage of the TALE on 4HBA, and their growth was compared to that of the parent strain, AW71, in M9 minimal medium with 15 mM 4HBA. Error shading indicates the standard deviation from the mean of three replicates, and assays were conducted in a microtiter plate format (BioscreenC).

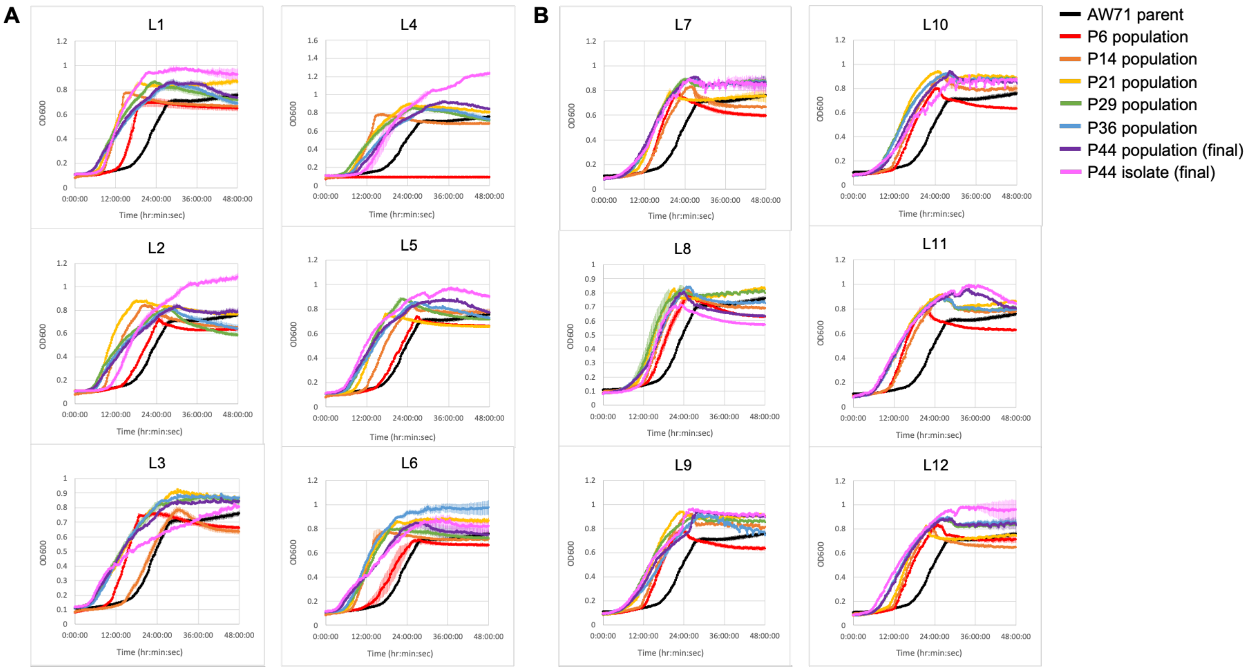

**Figure S3. Growth phenotype of populations sampled periodically throughout TALE, using 15 mM of aromatic substrate.** Samples of evolving populations were cryopreserved from each lineage (L1-L12) at different points during the TALE experiment (P6 = sixth passage, P14 = fourteenth passage, etc.). At the end of TALE, the growth of each sample was compared to the parent strain, AW71, and to population and isolate samples taken from the final passage (P44). Cultures were grown in microtiter plates (BioscreenC) in minimal medium with **(A)** 15 mM PCA, for lineages evolved on PCA, or **(B)** 15 mM 4HBA, for lineages evolved on 4HBA. Error shading indicates the standard deviation from the mean of three replicates.

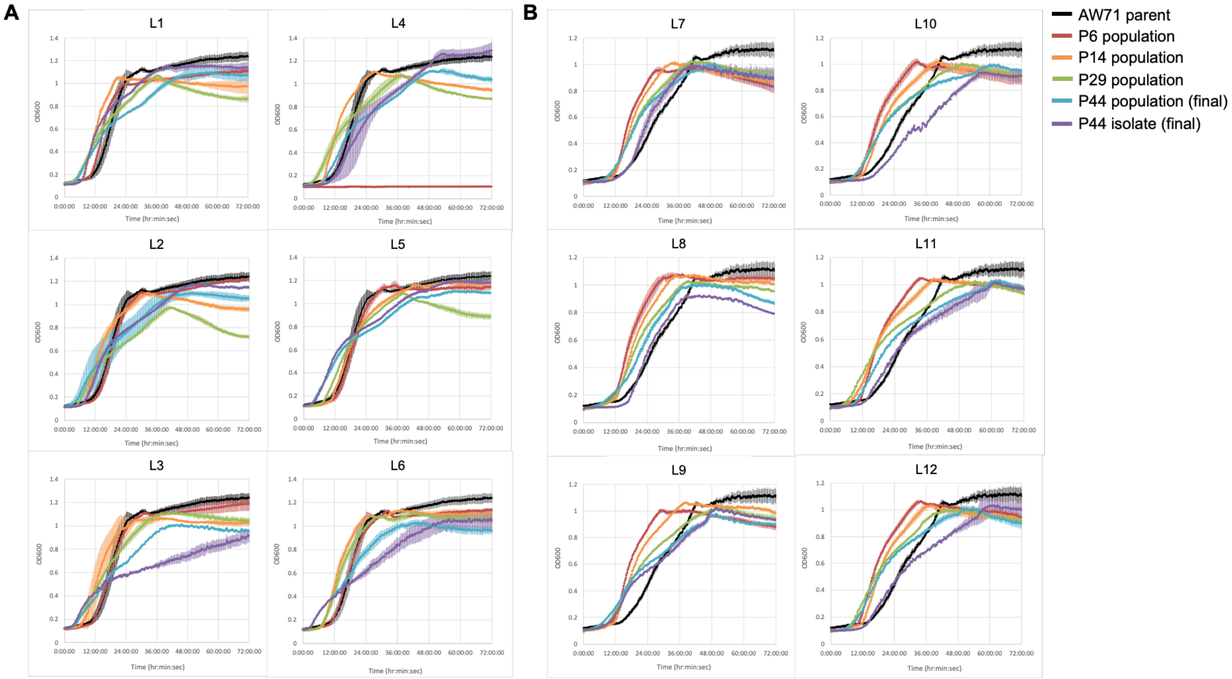

**Figure S4. Growth phenotype of populations sampled periodically throughout TALE, using 30 mM of aromatic substrate.** Samples of evolving populations were cryopreserved from each lineage (L1-L12) at different points during the TALE experiment (P6 = sixth passage, P14 = fourteenth passage, etc.). At the end of TALE, the growth of each sample was compared to the parent strain, AW71, and to population and isolate samples taken from the final passage (P44). Cultures were grown in microtiter plates (BioscreenC) in minimal medium with **(A)** 30 mM PCA, for lineages evolved on PCA, or **(B)** 30 mM 4HBA, for lineages evolved on 4HBA. Error shading indicates the standard deviation from the mean of three replicates.

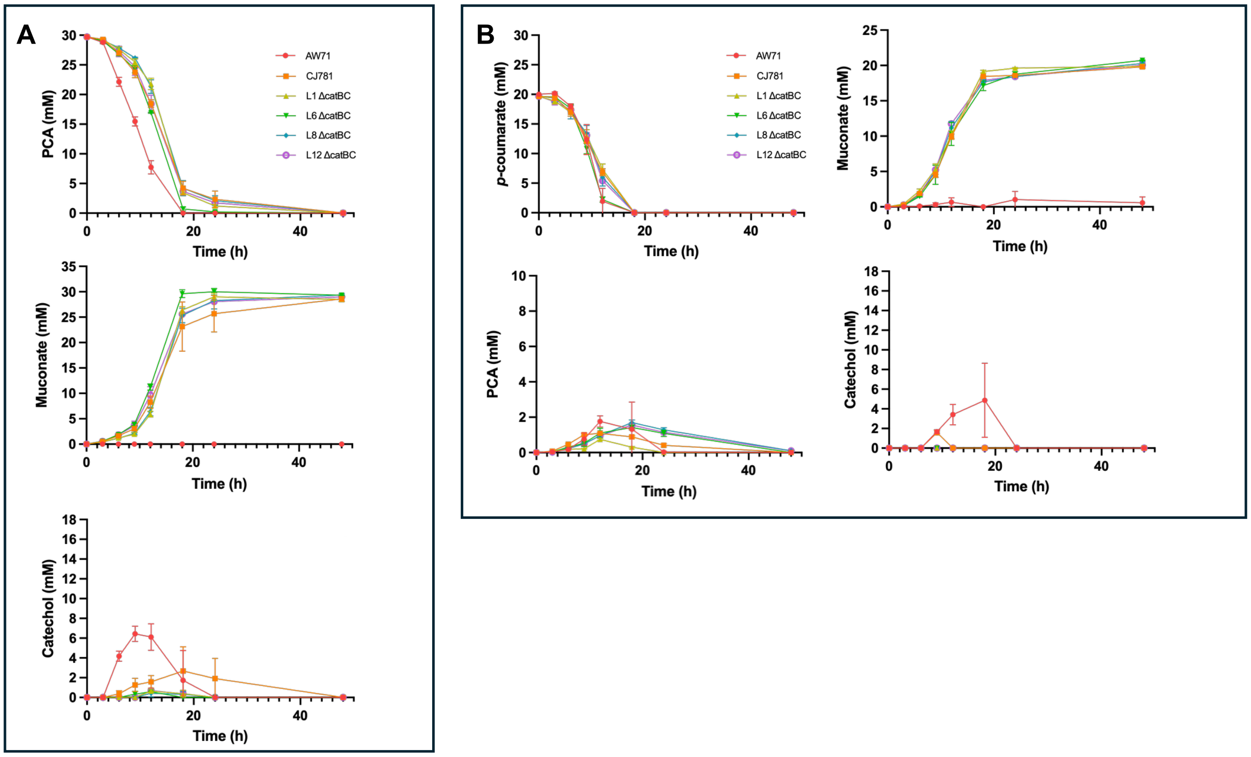

**Figure S5.** Isolates from TALE displayed decreased intermediate accumulation and improved muconate production upon deletion of *catBC*. The *catBC* genes were deleted in two evolved isolates per TALE condition (L1 and L6 from TALE on PCA; L8 and L12 from TALE on 4HBA). The resulting strains were cultivated in shaken flasks in M9 minimal medium with glucose to support growth and either **(A)** 30 mM PCA or **(B)** 20 mM *p*-coumarate. Error bars represent the standard deviation from the mean of three biological replicates.

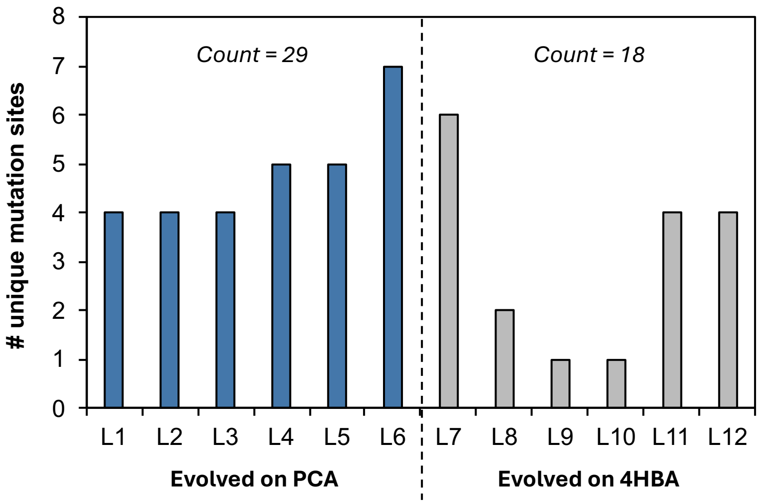

**Figure S6.** The number of unique mutation sites was tabulated for isolates evolved on PCA (blue) and isolates evolved on 4HBA (gray). Lineages evolved on PCA had a higher total mutation count than those evolved on 4HBA.

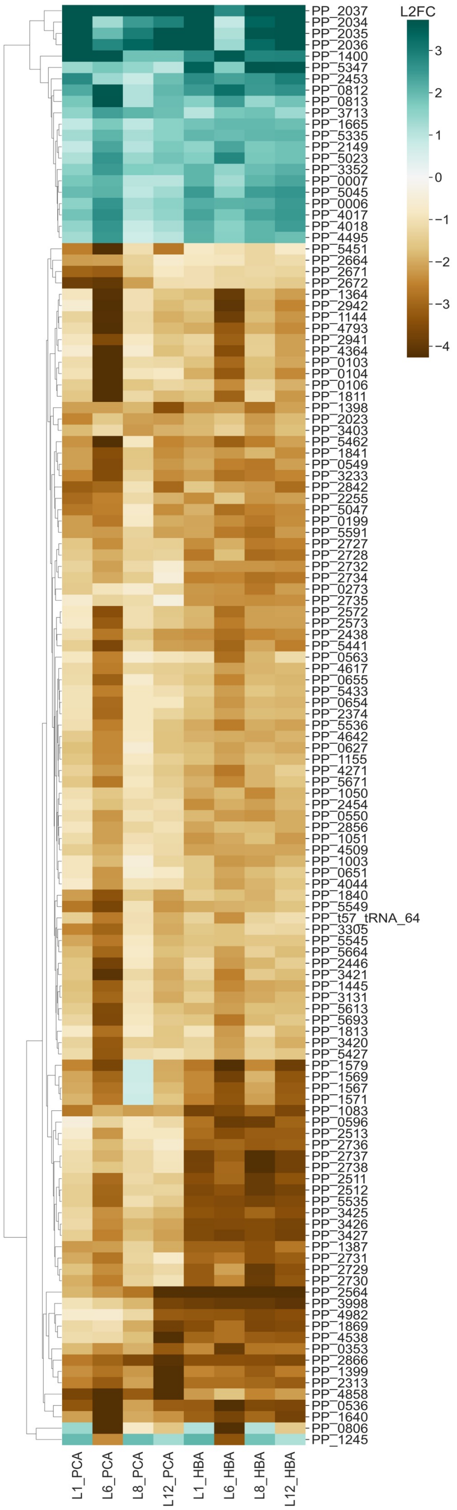

**Figure S7.** Heatmap of gene expression levels in evolved isolates, relative to the AW71 parent. Evolved isolates from L1, L6, L8, and L12 were cultivated in minimal medium with 30 mM PCA or 30 mM 4HBA, and cell pellets were collected at mid-log phase for transcriptomic analysis. Data is plotted only for genes where the magnitude of the $\log_2$(fold change) (L2FC) was >|2| in at least one condition.

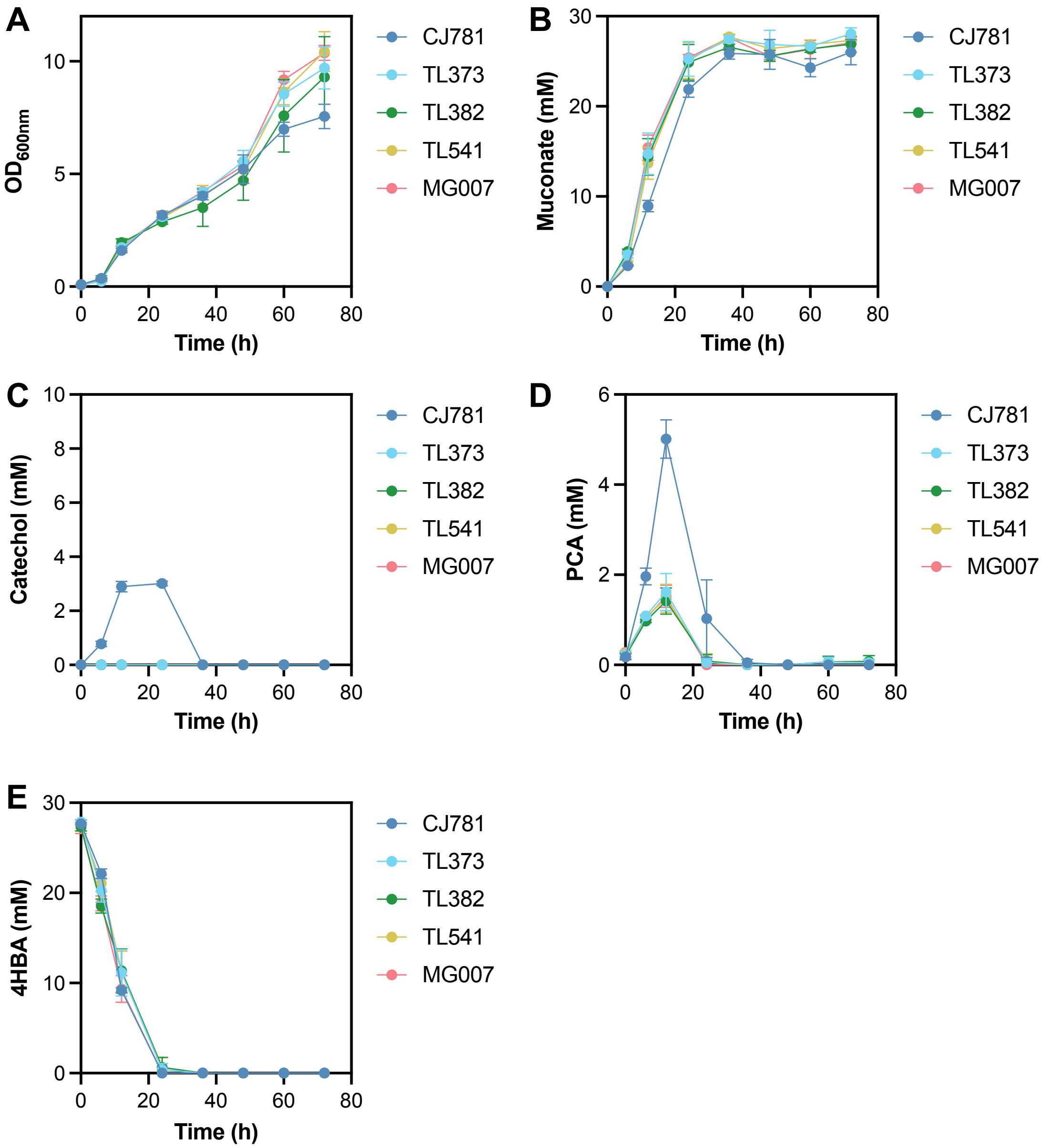

**Figure S8.** Reverse-engineering of ALE-derived mutations to the *catA1* cassette in the CJ781 strain background. Strains were cultivated in shake flasks in M9 minimal medium with 15 mM glucose + 30 mM 4HBA, and glucose was added to a concentration of 15 mM at 12 h and 24 h to support growth. The following metrics were tracked as a function of time: **(A)** cell density (optical density, OD, at 600nm), **(B)** muconate concentration, **(C)** catechol concentration, **(D)** PCA concentration, and **(E)** 4HBA concentration. Error bars indicate the standard deviation from the mean of three biological replicates.

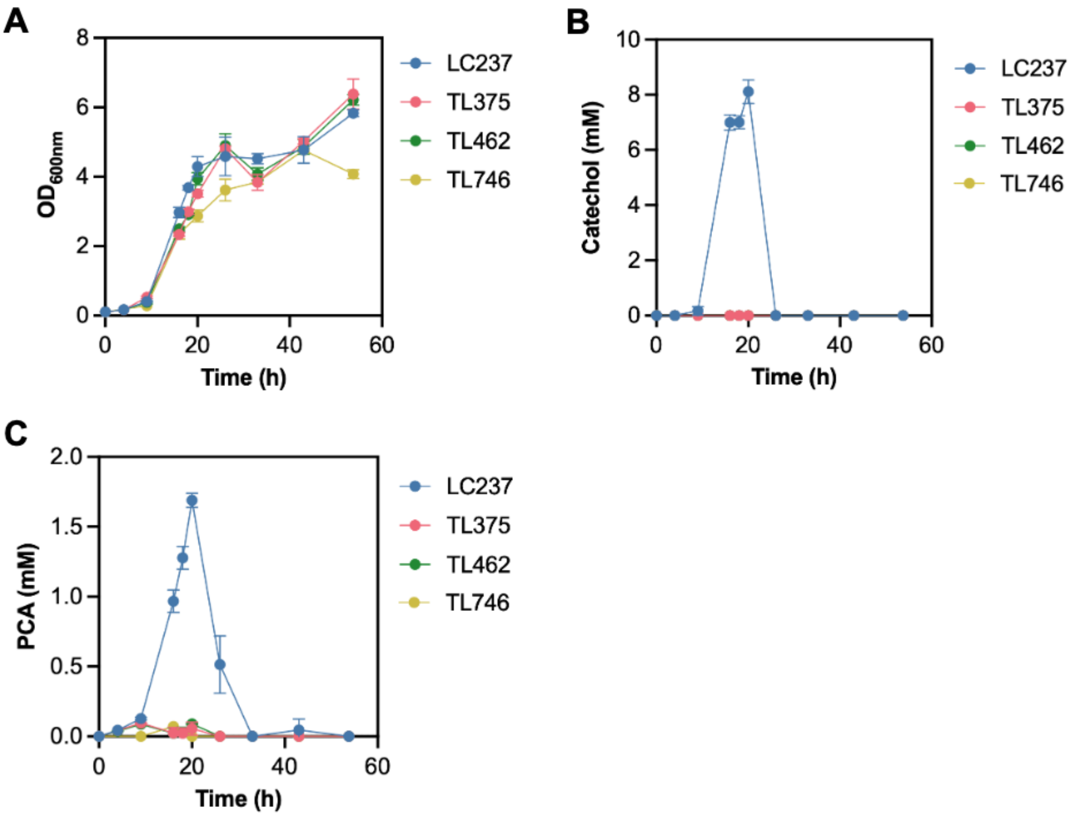

**Figure S9.** Reverse-engineering of ALE-derived mutations to the *catA1* cassette in the LC237 strain background. Strains were cultivated in shake flasks in M9 minimal medium with mock hydrolysate sugars. The following metrics were tracked as a function of time: **(A)** cell density (optical density, OD, at 600nm), **(B)** catechol (CAT) concentration, and **(C)** PCA concentration. Error bars indicate the standard deviation from the mean of three biological replicates.

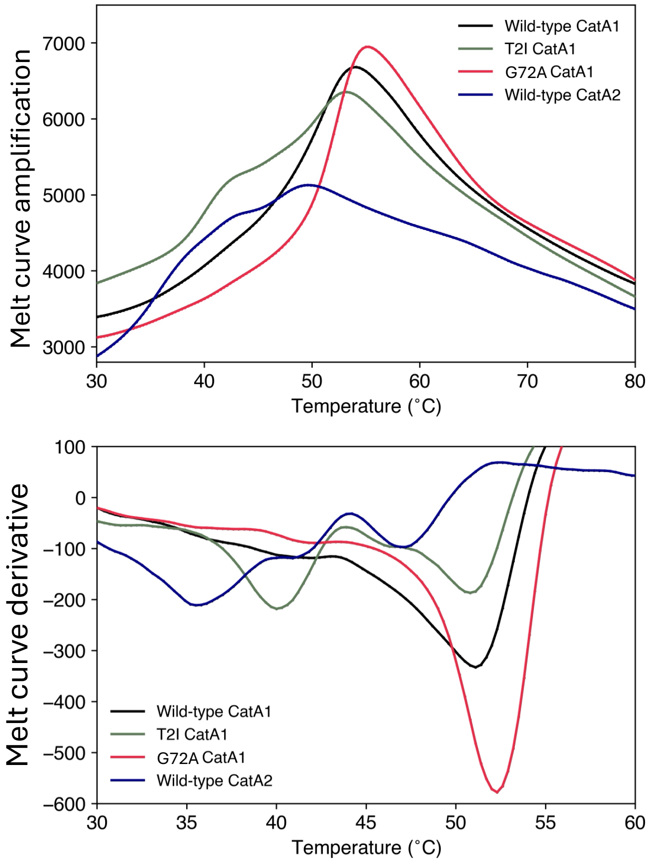

**Figure S10.** Melting temperature measurements for the catechol dioxygenases examined in this study. Differential scanning fluorimetry spectra (top) and its derivative (bottom) were used to determine the melting temperature of the different proteins. The assay was conducted in 20 mM HEPES, 100 mM NaCl, pH 7.5 and Sypro Orange Dye. The melting temperature was measured as the highest temperature where the derivative inflection is achieved.

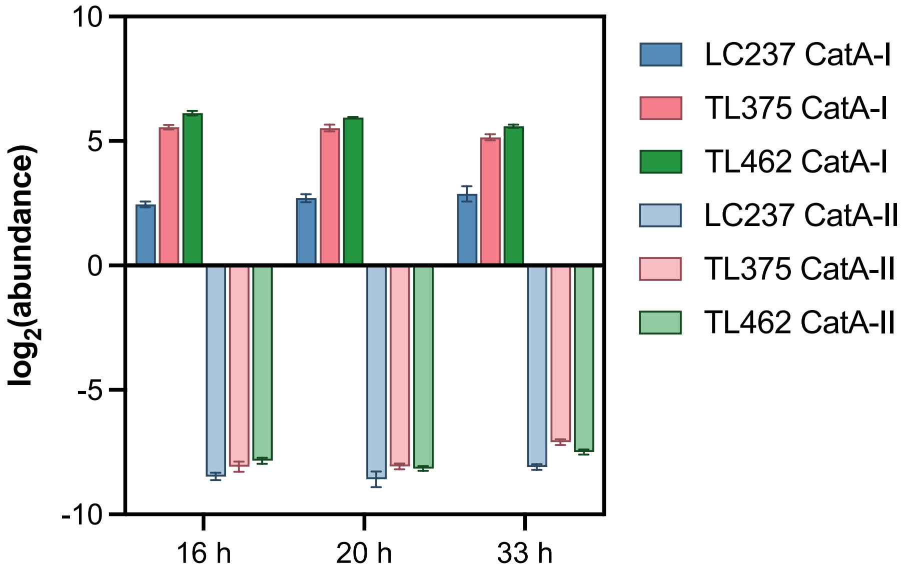

**Figure S11.** Quantitative proteomics determined the abundance of the CatA1 and CatA2 proteins in LC237, TL375, and TL462. Quantities are plotted as the $\log_2$ of abundance at each of the indicated timepoints during cultivation in shake flasks with M9 minimal medium and mock hydrolysate sugars. Error bars indicate the standard deviation from the mean of three biological replicates.

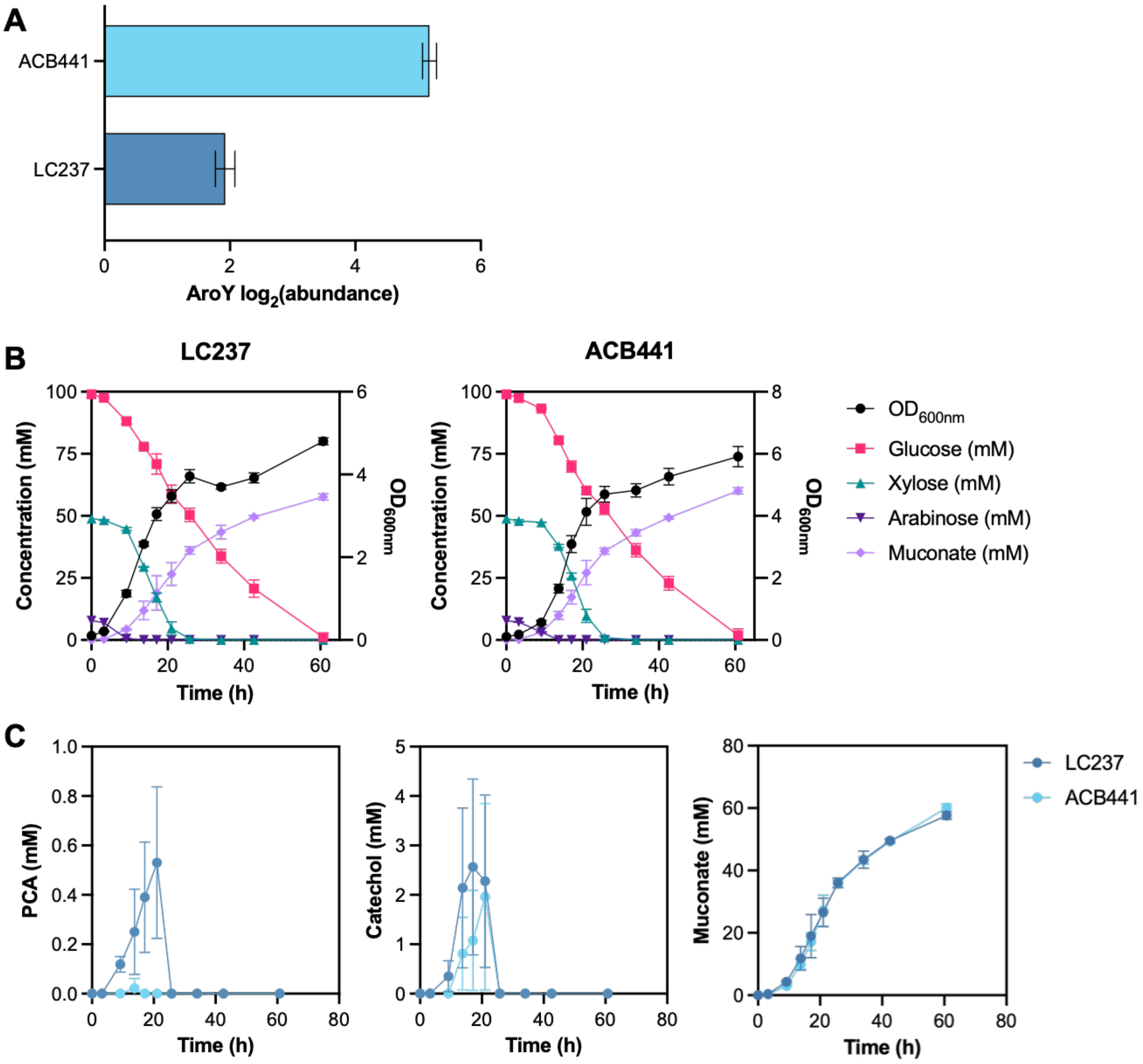

**Figure S12.** *P. putida* strain ACB441 contains a modified RBS sequence, designed to increase translation of AroY. **(A)** Quantitative proteomics showed that strain ACB441 exhibited increased AroY protein levels, relative to LC237, during **(B)** growth in shake flasks with M9 minimal medium and mock hydrolysate sugars. Error bars indicate the standard deviation from the mean of three biological replicates. **(B)** The accumulation of PCA was reduced in strain ACB441, compared to LC237, but catechol continued to accumulate as a metabolic intermediate, and no significant change in muconate level was observed between the two strains.

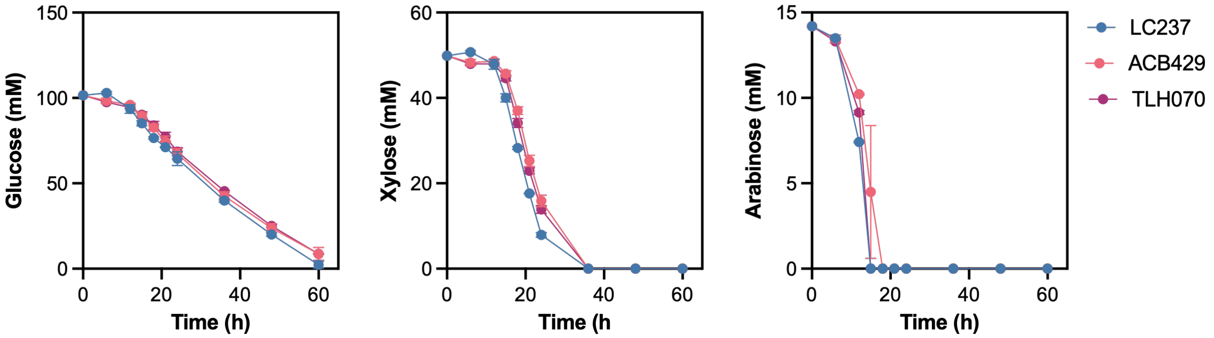

**Figure S13.** *P. putida* strains ACB429 and TLH070 did not show significant differences in sugar consumption, relative to LC237. Strains were cultivated in shake flasks with minimal medium containing 25 g/L mock hydrolysate. Error bars indicate the standard deviation from the mean of biological triplicates.

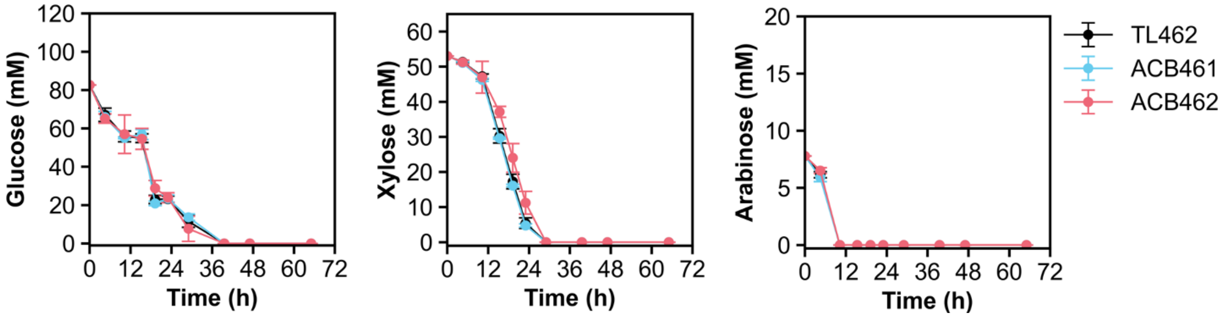

**Figure S14.** *P. putida* strains ACB461 and ACB462 did not show significant differences in substrate consumption, relative to LC237. Catechol was not detected at any timepoint in any strain, so it is not included in this figure. Strains were cultivated in shake flasks with minimal medium containing 25 g/L mock hydrolysate. Error bars indicate the standard deviation from the mean of three biological replicates.

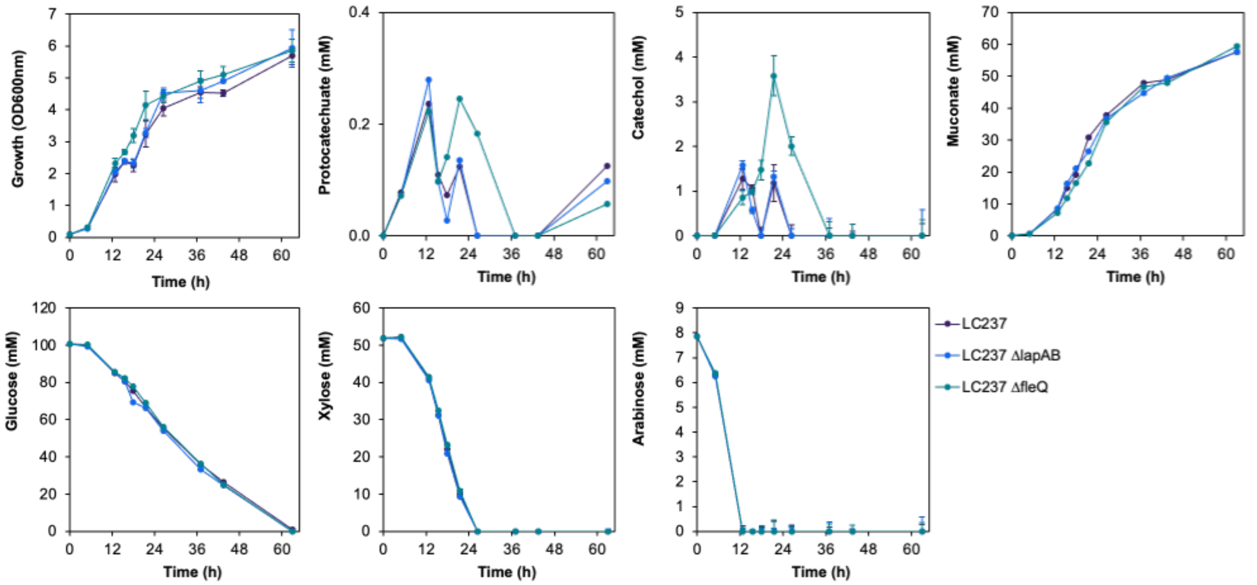

**Figure S15.** *P. putida* strains ACB458 (LC237 Δ*lapAB*) and ACB459 (LC237 Δ*fleQ*) did not show significant differences in intermediate accumulation, muconate production, or substrate consumption, relative to LC237. Strains were cultivated in shake flasks with minimal medium containing 25 g/L mock hydrolysate. Error bars indicate the standard deviation from the mean of three biological replicates.

**gnl|Q88GK8|Q88GK8_PSEPK|G18UU-21166** *catechol 1,2-dioxygenase, catA-I*
MTVK**ISHTADIQAFFNRVAGLDHAEGNPR**FKQIILR**VLQDTARLIEDLEITEDEFWHAVDYLNR**LGGR**NEAGLLAAGLGIEHF**
**LDLLQDAKDAEAGLGGGTPRTIEGPLYVAGAPLAQGEAR**MDDGTDPGVVMFLQGQVFDADGKPLAGATVDLWHANTQGTY
SYFDSTQSEFNLRRR**IITDAEGR**YRAR**SIVPSGYGCDPQGPTQECLDLLGRHGQRPAHVHFFISAPGHRHLTTQINFAGDKY**
**LWDDFAYATRDGLIGELRFVEDAAAAR**DR**GVQGERFAELSFDFR**LQGAK**SPDAEAR**SHRPRALQEG

**gnl|Q88I35|Q88I35_PSEPK|G18UU-20592** *catechol 1,2-dioxygenase, catA-II*
MTVNISHTAEVQQFFEQAAGFCNAAGNPRLKRIVQR**LLQDTARLIEDLDISEDEFWHAVDYLNR**LGGR**GEAGLLVAGLGIEH**
**FLDLLQDAKDQEAGRVGGTPRTIEGPLYVAGAPIAQGEVR**MDDGSEEGVATVMFLEGQVLDPHGRPLPGATVDLWHANTR
**GTYSFFDQSQSAYNLR**RR**IVTDAQGR**YRAR**SIVPSGYGCDPQGPTQECLDLLGRHGQRPAHVHFFISAPGYRHLTTQINLS**
**GDKYLWDDFAYATRDGLVGEVVFVEGPDGR**HAELK**FDFQLQQAQGGADEQR**SGRPRALQEA

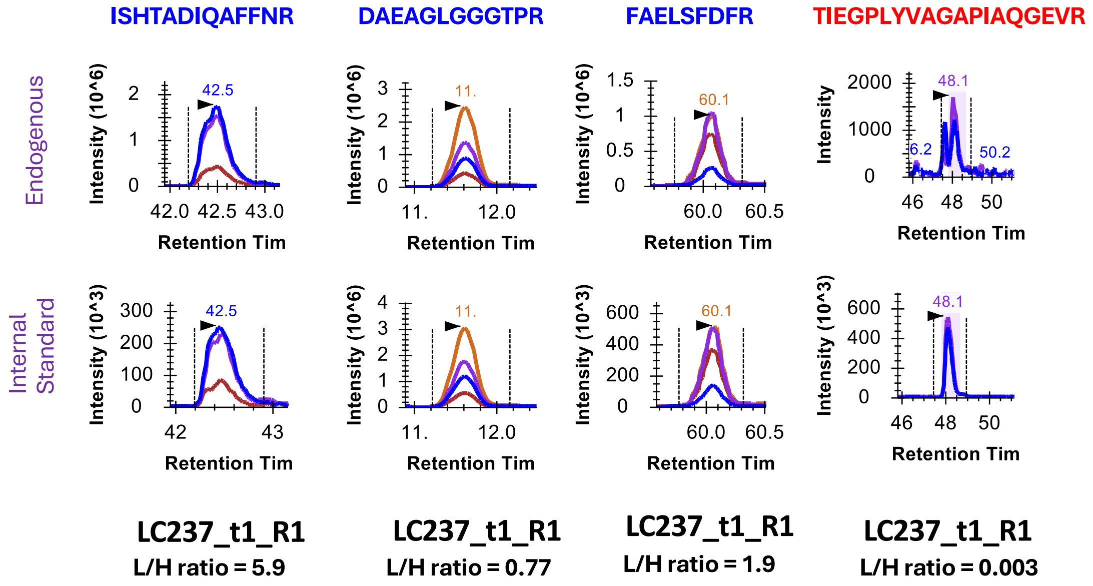

**Figure S16.** Peptide selection and LC peaks for CatA1 and CatA2 targeted proteomics. The full sequence of each protein is shown at top, with theoretically detectable peptide sequences shown in bold. Peptide sequences selected for detection of CatA1 are shown in blue text, and the peptide sequence used for CatA2 is shown in red text. Plots indicate peak area ratios between endogenous light peptides (L) and their corresponding heavy isotope-labeled internal standards (H) for a representative sample (strain LC237 sampled at t1 = 16 h, R1 = replicate 1).

**Table S1.** Construction details for engineered bacteria generated in this study.

| Strain | Genotype | Construction details[a] |
|---|---|---|
| **KT2440** | *Pseudomonas putida* KT2440 | Wild-type strain, ATCC 47054. $Cm^r$ $Amp^r$ |
| **Evolved isolates ∆*catBC*** | Evolved isolates L1, L6, L8, and L12, each with ∆*catBC* modification | Evolved isolates were transformed with 100 ng each of pCJ033. Colonies were selected twice on LB + 50 mg/L kanamycin agar and twice on YT + 25% sucrose agar, and then colony PCR with oCJ261/oCJ262 was performed to identify *catBC* deletion mutants. |
| **LC237** | *P. putida* KT2440 Δ*catRBC*::$P_{tac}$:*catA1* Δ*pcaHG*::$P_{tac}$:*aroY*:*ecdB*:*asbF* Δ*pykA*::*aroG-D146N*:*aroY*:*ecdB*:*asbF* Δ*pykF* Δ*ppc* Δ*pgi-2* Δ*gcd* Δ*hexR* Δ*ampC*::$P_{xylE*}$*xylE*:P*tac*:*xylAB*:*talB*:*tktA* PP_1736-1737(intergenic)::$P_{lac}$:*ubiC*-C22 *xylE-A62V, A455V* $P_{PP\_2569}$ G→A Δ*pykF*::$P_{tac}$:*aroB* ∆*gcd*::*araE-araCDABE* | Kim *et al.*[1] |
| **TL375** | LC237 *catA1*$_{T2I}$ | LC237 was transformed with 100 ng of pTL068. Colonies were selected twice on LB + 50 mg/L kanamycin agar and twice on YT + 25% sucrose agar, and then colony PCR with oLC320/oLC080 was performed to identify a clone with the correct T2I mutation based on Sanger sequencing. |
| **TL462** | LC237 *catA1* RBS C→G | LC237 was transformed with 100 ng of pTL070. Colonies were selected twice on LB + 50 mg/L kanamycin agar and twice on YT + 25% sucrose agar, and then colony PCR with oLC320/oLC080 was performed to identify a clone with the correct RBS C>G mutation based on Sanger sequencing. |
| **TL746** | LC237 *catA1* RBS C→G, *catA1*$_{G72A}$ | LC237 was transformed with 100 ng of pTL131. Colonies were selected twice on LB + 50 mg/L kanamycin agar and twice on YT + 25% sucrose agar, and then colony PCR with oLC320/oLC080 was performed to identify a clone with the correct RBS C>G, G72A mutation based on Sanger sequencing. |
| **TLH070** | LC237 PP_2553-2554 intergenic A→C | LC237 was transformed with 100 ng of pTLH011. Colonies were selected twice on LB + 50 mg/L kanamycin agar and twice on YT + 25% sucrose agar, and then colony PCR with oCJ680/oCJ681 was performed to identify a clone with the correct intergenic mutation based on Sanger sequencing. |
| **CJ781** | *P. putida* Δ*catRBCA*::$P_{tac}$:*catA1* Δ*pcaHG*::$P_{tac}$:*aroY:ecdBD* *fpvA*:$P_{tac}$:*praI:vanAB* Δ*pobAR* Δ*crc* | Kuatsjah *et al.*[2] |
| **AW071** | *P. putida* Δ*catRBCA*::$P_{tac}$:*catA1* Δ*pcaHG*::$P_{tac}$:*aroY:ecdBD* *fpvA*:$P_{tac}$:*praI:vanAB* Δ*pobAR* Δ*crc*::$P_{tac}$:*catB:catC* | CJ781 was transformed with pAW022. Colonies were selected twice on LB + 50 mg/L kanamycin agar and twice on TY + 25% sucrose agar. The strain was confirmed by colony PCR with oCJ261 and oCJ262 (3605 bp, $T_m$=65°C) and sequencing. |

| | | |
|---|---|---|
| **MG007** | CJ781 *catA1* RBS C→G, *catA1*$_{T2I,G72A}$ | CJ781 was transformed with pMG005. Colonies were selected twice on LB + 50 mg/L kanamycin agar and twice on TY + 25% sucrose agar. Colony PCR with primers oLC320/oLC080 was used to amplify the region of interest. Nanopore sequencing of the PCR products was used to sequence verify the strain. |
| **TL373** | CJ781 *catA1*$_{T2I}$ | CJ781 was transformed with 100 ng of pTL068. Colonies were selected twice on LB + 50 mg/L kanamycin agar and twice on YT + 25% sucrose agar, and then colony PCR with oLC320/oLC080 was performed to identify a clone with the correct T2I mutation based on Sanger sequencing. |
| **TL382** | CJ781 *catA1* RBS C→G | CJ781 was transformed with 100 ng of pTL070. Colonies were selected twice on LB + 50 mg/L kanamycin agar and twice on YT + 25% sucrose agar, and then colony PCR with oLC320/oLC080 was performed to identify a clone with the correct RBS C>G mutation based on Sanger sequencing. |
| **TL541** | CJ781 *catA1* RBS C→G, *catA1*$_{G72A}$ | CJ781 was transformed with 100 ng of pTL131. Colonies were selected twice on LB + 50 mg/L kanamycin agar and twice on YT + 25% sucrose agar, and then colony PCR with oLC320/oLC080 was performed to identify a clone with the correct RBS C>G, G72A mutation based on Sanger sequencing. |
| **ACB429** | LC237 ΔPP_5042::P$_{lac}$:*dxs*:*dxr* | LC237 was transformed with 1000 ng of pACB169. Colonies were selected twice on LB + 50 mg/L kanamycin agar and twice on YT + 25% sucrose agar. Colony PCR with primers oACB466/oACB467 was used to amplify the region of interest. Nanopore sequencing of the PCR product was used to verify integration of the *dxs*:*dxr* cassette. |
| **ACB441** | LC237 *aroY* strengthened RBS | LC237 was transformed with 1000 ng of pRW61. Colonies were selected twice on LB + 50 mg/L kanamycin agar and twice on YT + 25% sucrose agar. Colony PCR with primers oCJ178/oLC270 was used to amplify the region of interest. Nanopore sequencing of the PCR product was used to verify integration of the modified *aroY* RBS sequence (AAAAAACCTCCTTAGGTTGAGTTTAGACTTCTTGG TCGCGGCCAATGGCC). |
| **ACB458** | LC237 Δ*lapAB* | LC237 was transformed with 1000 ng of pCA010. Colonies were selected twice on LB + 50 mg/L kanamycin agar and twice on YT + 25% sucrose agar. Colony PCR with primers oCA051/oCA052 was used to amplify the region of interest. Nanopore sequencing of the PCR product was used to verify *lapAB* deletion. |
| **ACB459** | LC237 Δ*fleQ* | LC237 was transformed with 1000 ng of pGB028. Colonies were selected twice on LB + 50 mg/L kanamycin agar and twice on YT + 25% sucrose agar. Colony PCR with primers oGB181/oGB182 was used to amplify the region of interest. Nanopore sequencing of the PCR product was used to verify *fleQ* deletion. |
| **ACB461** | TL462 Δ*lapAB* | TL462 was transformed with 1000 ng of pCA010. Colonies were selected twice on LB + 50 mg/L |

| | | |
|---|---|---|
| | | kanamycin agar and twice on YT + 25% sucrose agar. Colony PCR with primers oCA051/oCA052 was used to amplify the region of interest. Nanopore sequencing of the PCR product was used to verify *lapAB* deletion. |
| **ACB462** | TL462 ∆*fleQ* | TL462 was transformed with 1000 ng of pGB028. Colonies were selected twice on LB + 50 mg/L kanamycin agar and twice on YT + 25% sucrose agar. Colony PCR with primers oGB181/oGB182 was used to amplify the region of interest. Nanopore sequencing of the PCR product was used to verify *fleQ* deletion. |

**Table S2.** Plasmids used in this study.

| Plasmid | Description[a] | Construction details |
|---|---|---|
| **pK18sB** | Sucrose counter-selection allelic exchange vector for *P. putida* KT2440; Km$^r$ | Previously described in Jayakody *et al.*[3] GenBank Accession #MH166772. Addgene Plasmid # 177838. |
| **pK18msB** | Sucrose counter-selection allelic exchange vector for *P. putida* KT2440 with mobilization factor required for conjugation; Km$^r$ | Previously described in Ling *et al.*[4] GenBank Accession #OK423783. Addgene Plasmid #177839. |
| **pAW022** | pk18sB-based plasmid for deletion of *crc* with simultaneous overexpression of *catBC* | Plasmid was synthesized by Twist Biosciences to include 1000 bp and 967 bp homology regions aligned immediately upstream and downstream of *crc* for deletion by homologous recombination. The *tac* promoter (ttgacaattaatcatcggctcgtataatgtgtggaattgtgagcg gataacaatttcacac) was used to drive expression of *catBC* in a two-gene operon. Synthetic RBSs were designed for *catB* (AACCTGGGGGCTTATTGACTACTATAAACA TTTTGCGGGGTATTAAA) and *catC* (GCGTTACCGCTGTATCCAAAGGAGATTTG TCG) using the Salis RBS Calculator [ref]. The *tonB* terminator (AGTCAAAAGCCTCCGACCGGAGGCTTTTT ACT) was placed at the end of the expression cassette. |
| **pTL068** | pK18sB-based plasmid for introduction of T2I mutation to *catA1* (PP_3713) in *P. putida* KT2440; Km$^r$ | Site directed mutagenesis was performed on pAW077 using oTL292/oTL293 to introduce T2I mutation. Linear fragment was ligated using KLD reaction mix and the resulting plasmid was Nanopore sequenced for verification of correct construct. |
| **pTL069** | pK18sB-based plasmid for introduction of G72A mutation to *catA1* (PP_3713) in *P. putida* KT2440; Km$^r$ | Site directed mutagenesis was performed on pAW077 using oTL294/oTL295 to introduce G72A mutation. Linear fragment was ligated using KLD reaction mix and the resulting plasmid was Nanopore sequenced for verification of correct construct. |
| **pTL070** | pK18sB-based plasmid for introduction of RBS C>G mutation to *catA1* (PP_3713) in *P. putida* KT2440; Km$^r$ | Site directed mutagenesis was performed on pAW077 using oTL296/oTL297 to introduce RBS C>G mutation. Linear fragment was ligated using KLD reaction mix and the resulting plasmid was Nanopore sequenced for verification of correct construct. |
| **pTL131** | pK18sB-based plasmid for introduction of RBS C>G, G72A mutation to *catA1* (PP_3713) in *P. putida* KT2440; Km$^r$ | Site directed mutagenesis was performed on pTL069 using oTL294/oTL295 to introduce G72A mutation. Linear fragment was ligated using KLD reaction mix and the resulting plasmid was Nanopore sequenced for verification of correct construct. |
| **pTLH011** | pK18msB-based plasmid for integration of mutation between | Linear fragment containing the target sequence and mutation (Twist bioscience) was amplified using oTLH067/oTLH068 and ligated using |

| | | |
|---|---|---|
| | PP_2553-2554 in *P. putida* KT2440; Km$^r$ | KLD reaction mix. The resulting plasmid was Nanopore sequenced for verification of correct construct. |
| **pEE083** | pET-based heterologous production of WT CatA1 (PP_3713) in *E. coli*; Am$^r$ | Kijpornyongpan *et al.*[5] |
| **pMG005** | pK18sB-based plasmid for introduction of RBS C>G, T2I, and G72A mutations to *catA1* (PP_3713) in *P. putida* KT2440; Km$^r$ | pMG005 was constructed by PCR amplifying pTL131 with oTL292/oTL593, treating the PCR product with NEB KLD mix, and transforming into NEB 5-alpha F'Iq via heat shock transformation according to NEB's protocol and plated on LB+Km50 plates. Plasmids were sequenced via Nanopore sequencing to verify the correct construct. |
| **pTL082** | pET-based heterologous production of G72A CatA1 (PP_3713) in *E. coli*; Am$^r$ | Subcloned by TL via Gibson assembly using pEE083 for vector template. |
| **pTL083** | pET-based heterologous production of WT CatA2 (PP_3166) in *E. coli*; Am$^r$ | Subcloned by TL via Gibson assembly using pEE083 for vector template. |
| **pRW61** | pK18sB-based plasmid for introduction of the stronger RBS on *aroY* | Wilkes *et al.*[6] |
| **pACB169** | pK18sB-based plasmid for overexpression of $P_{lac}$:*dxs*:*dxr* at the PP_5042 locus. | Fragments for *dxs* (PP_0527) and *dxr* (PP_1597) were amplified from *P. putida* KT2440 gDNA using primer pairs oACB607/oACB608 and oACB609/oACB610, respectively, which added a synthetic RBS on each CDS as well as a *lac* promoter at the beginning of the synthetic operon. pACB124, a pK18sB-based plasmid for integration at the PP_5042 locus, was amplified with primers oACB611/oACB612. The three fragments were assembled by the method of Gibson in a 1:2:2 (plasmid:*dxs*:*dxr*) molar ratio and the reaction was transformed into NEB 5-alpha F'Iq *E. coli* cells. Colonies were selected on LB+Km50 plates, and plasmids were sequenced by Nanopore sequencing to verify the correct construct. |
| **pCA010** | pK18sB-based plasmid for deletion of *lapAB* (PP_0167-0168) | Borchert *et al.*[7] |
| **pGB028** | pk18sB-based plasmid for deletion of *fleQ* (PP_4373) | Borchert *et al.*[7] |
| **pCJ033** | pk18sB-based plasmid for deletion of the *crc* (PP_5292) locus (used to delete *catBC* in evolved isolates of AW071) | Johnson *et al.*[8] |

**Table S3.** DNA Sequences of oligos used in this study. Integrated DNA Technologies (IDT) was used for synthesis.

| Primer | Sequence (5'→3') | Description |
|---|---|---|
| **oTL292** | AGCACGATGATCGTGAAAAT TTC | Mutagenesis: paired with oTL293; introduces T2I mutation to *catA1* (PP_3713) in pTL068. |
| **oTL293** | GACCTCGTATTGTGTGAAAT TG | Mutagenesis: paired with oTL292; introduces T2I mutation to *catA1* (PP_3713) in pTL068. |
| **oTL294** | GCTGCCTCGTTACGGCCGC C | Mutagenesis: paired with oTL295; introduces G72A mutation to *catA1* (PP_3713) in pTL069 and pTL131. |
| **oTL295** | CCTGCTGGCTGCTGGCCTG G | Mutagenesis: paired with oTL294 to introduce G72A mutation to *catA1* (PP_3713) in pTL069 and pTL131. |
| **oTL296** | TCACACAATAGGAGGTCAGC AC | Mutagenesis: paired with oTL297 to introduce RBS C>G mutation to *catA1* (PP_3713) in pTL070. |
| **oTL297** | AATTGTTATCCGCTCACAATT C | Mutagenesis: paired with oTL296 to introduce RBS C>G mutation to *catA1* (PP_3713) in pTL070. |
| **oTL593** | TCATCGTGCTGACCTCCTAT TG | For amplification of pTL131 with oTL292, and introduction of the C>G SNP in the *catA1* RBS. |
| **oLC270** | CACCCGCTGCGCGGAAGT | Diagnostic/sequencing: binds adjacent to *pcaHG* site in *P. putida*. |
| **oLC080** | AAGCTGTCGCTGAGCCTGTT | Diagnostic/sequencing: binds upstream of the *catA1* gene (PP_ PP_3713) to check sequence of *catA1* mutations in *P. putida*. |
| **oLC320** | CCTGAACCTTCGAAGCGCAT | Diagnostic/sequencing: binds downstream of the *catA1* gene (PP_ PP_3713) to check sequence of catA mutations in *P. putida*. |
| **oTLH067** | TGACATGATTACGAATTCAA GGCCGGTGCGAGGTAG | Amplification: paired with oTLH068 to generate pTLH011, reverse engineering mutations between PP_2553-2554 in *P. putida.* |
| **oTLH068** | ACGGCCAGTGCCAAGCTTTT CGCCCAGCAGGCGC | Amplification: paired with oTLH067 to generate pTLH011, reverse engineering mutations between PP_2553-2554 in *P. putida.* |
| **oCJ178** | TCACTTCTTGTCGCTGAACA GCTCTGG | Diagnostic/sequencing: reverse primer for *aroY* CDS. |
| **oCJ261** | GCCATGAATAGCTGCTCC | Colony PCR to confirm integration of pAW022. |
| **oCJ262** | GGTCAGCGTTGAAAACGG | |
| **oCJ680** | TTTGTGATGCTCGTCAGGG | Diagnostic/sequencing: binds upstream of the insert to check sequence of mutations in *P. putida*. |
| **oCJ681** | CTTCCCAACCTTACCAGAG | Diagnostic/sequencing: binds downstream of the insert to check sequence of mutations in *P. putida*. |

| | | |
|---|---|---|
| **oACB466** | CAGGTTGCTCAGGCTGTCGTAC | Diagnostic/sequencing: binds upstream of PP_5042 in *P. putida*. |
| **oACB467** | GCATTGGCTGCGTAACGAAGC | Diagnostic/sequencing: binds downstream of PP_5042 in *P. putida*. |
| **oACB607** | GTTGTGTGGAATTGTGAGCGGATAACAATTTCACACCACTACCCCAGAACCTAGGAAATAGAACATGCCCACGACGTTTCAAGAG | With oACB608, amplifies *dxs* from KT2440 gDNA for assembly of pACB169. Fwd |
| **oACB608** | CCTTCTCCGCTATTGCACGCGAAAGGACCCTAGAGC | With oACB607, amplifies *dxs* from KT2440 gDNA for assembly of pACB169. Rev |
| **oACB609** | TGCAATAGCGGAGAAGGAAATTTTAATGGGGTGTGATGTGAGTCG | With oACB610, amplifies *dxr* from KT2440 gDNA for assembly of pACB169. Fwd |
| **oACB610** | CTTCACAGGTCCAGCACCAGTTGGCGGGCCTGCATCAG | With oACB609, amplifies *dxr* from KT2440 gDNA for assembly of pACB169. Rev |
| **oACB611** | GCTCACAATTCCACACAACATACGAGCCGGAAGCATAAAGTGTAAAGCCTGGGGTGCCTAATGCAACCACCTTGGGCTTGTAGGC | With oACB612, amplifies backbone for assembly of pACB169. Fwd |
| **oACB612** | ATGCAGGCCCGCCAACTGGTGCTGGACCTGTGAAG | With oACB611, amplifies backbone for assembly of pACB169. Rev |
| **oCA051** | TCAATGACCAGCGTGTCGT | Diagnostic/sequencing: binds upstream of *lapAB* in *P. putida*. |
| **oCA052** | TGGTCTGTCAGCTGTCCTT | Diagnostic/sequencing: binds downstream of *lapAB* in *P. putida*. |
| **oGB181** | GGATACCCTCGTCGCCAAGC | Diagnostic/sequencing: binds upstream of *fleQ* in *P. putida*. |
| **oGB182** | GCTCACCCAGATGACTGTCGC | Diagnostic/sequencing: binds downstream of *fleQ* in *P. putida*. |
